# Low spontaneous mutation rate and Pleistocene radiation of pea aphids

**DOI:** 10.1101/769133

**Authors:** Varvara Fazalova, Bruno Nevado

## Abstract

Accurate estimates of divergence times are essential to understand the evolutionary history of species. It allows linking evolutionary histories of the diverging lineages with past geological, climatic and other changes in environment and shed light on the processes involved in speciation. The pea aphid radiation includes multiple host races adapted to different legume host plants. It is thought that diversification in this system occurred very recently, over the past 8,000 to 16,000 years. This young age estimate was used to link diversification in pea aphids to the onset of human agriculture, and lead to the establishment of the pea aphid radiation as a model system in the study of speciation with gene flow. Here, we re-examine the age of the pea aphid radiation, by combining a mutation accumulation experiment with a genome-wide estimate of divergence between distantly related pea aphid host races. We estimate the spontaneous mutation rate for pea aphids as 2.27 × 10^−10^ per haploid genome per parthenogenic generation. Using this estimate of mutation rate and the genome-wide genetic differentiation observed between pea aphid host races, we show that the pea aphid radiation is much more ancient than assumed previously, predating Neolithic agriculture by several hundreds of thousands of years. Our results rule out human agriculture as the driver of diversification of the pea aphid radiation, and call for re-assessment of the role of allopatric isolation during Pleistocene climatic oscillations in divergence of the pea aphid complex.

## Introduction

The pea aphid (*Acyrthosiphon pisum*, Harris) radiation includes at least 15 host races (Peccoud et al. 2015) specialized on different legume host plants (e.g., Müller 1962; Via 1991; Ferrari et al. 2006) that represent genetically divergent lineages (e.g., Peccoud et al. 2009a; Ferrari et al. 2012). Previous work suggests that diversification in this host race complex occurred very rapidly, i.e. within the past 8,000 – 16,000 years ago (Peccoud et al. 2009b). This timing of divergence was used to link diversification of pea aphid host races to the increased availability of potential host plants due to climate warming and the onset of Neolithic agriculture (Peccoud et al. 2009b). Hybrids between most host races are relatively common in nature and this – together with the assumed recent divergence – has been interpreted as evidence for a scenario of divergence with gene flow in the pea aphid complex. Pea aphids have since been established as a model system for studying speciation with gene flow (Peccoud and Simon 2010).

The pea aphid radiation includes a few host races that do not hybridise in nature (Peccoud et al. 2009a), and intrinsic reproductive isolation between some races has recently been shown (Fazalova et al. 2018). This very rapid completion of speciation is at odds with a scenario of very recent divergence with gene flow, and indeed contrasts with other insect taxa for which the build-up of intrinsic reproductive isolation takes much longer, e.g. at least 1,500,000 years in *Heliconius* butterflies (Kozak et al. 2015) or over 200,000 years in *Drosophila* flies (Coyne and Orr 1997). Accounting for different generation times in these groups, this would suggest that completion of speciation takes over 6 million generations in *Heliconius* (4 generations per year, (Keightley et al. 2015)) and over 3 million generations in *Drosophila* (15 generations per year, (Pool 2015)) but only 120-240 thousand generations in pea aphids (15 generations per year, (Loxdale and Balog 2018)).

The mismatch between time taken to achieve strong intrinsic reproductive isolation in pea aphids compared to other insect taxa calls for re-examination of the age of this complex. To do so, we estimated genome-wide spontaneous mutation rate for several pea aphid host races with a mutation accumulation experiment; and estimated average genomic differentiation for two distantly related host races. Together, these analyses show that the pea aphid radiation is much older than previously assumed, at least 543,000 years (95% CI 419,000 – 772,000). This new estimate implies that the time needed for completion of speciation in pea aphids is similar to that estimated in other insect systems, and calls for re-interpretation of the mechanisms driving diversification in pea aphids – in particular, the role of geographic isolation in the diversification of the pea aphid radiation needs to be reassessed.

## Results and Discussion

### Low spontaneous mutation rate in pea aphids

In order to obtain an estimate of the spontaneous mutation rate of the pea aphid, we performed a mutation accumulation experiment, with twelve parthenogenetic lines (representing four host races) over 28 generations. For each mutation accumulation line, we re-sequenced genomes to a mean coverage 38×, trimmed and mapped sequence data to the pea aphid genome, and identified de novo mutations using a strict bioinformatics pipeline (see Material and Methods). On average 91% of reads were mapped to the reference genome assembly (Table S1) and on average 44% of the genome was callable within each line (i.e. had called genotypes at both the start and end generations; Table 1).

**Table 1.** Number of callable sites, number of mutations and mutation rate (uncorrected for false positives) for each asexual lineage.

| Line | Host race | Number of callable sites | Number of mutations | Mutation rate (per site per haploid genome per generation) |
| --- | --- | --- | --- | --- |
| 05b | *M. lupulina* | 175,281,226 | 4 | $4.2 \times 10^{-10}$ |
| 09b | *L. corniculatus* | 185,010,536 | 5 | $5.0 \times 10^{-10}$ |
| 107a | *Lat. pratensis* | 196,589,460 | 2 | $1.9 \times 10^{-10}$ |
| 118a | *L. corniculatus* | 280,177,299 | 4 | $2.6 \times 10^{-10}$ |
| 120a | *Lat. pratensis* | 189,053,054 | 8 | $7.8 \times 10^{-10}$ |
| 122a | *L. corniculatus* | 258,621,867 | 1 | $7.2 \times 10^{-11}$ |
| 29d | *Lat. pratensis* | 155,191,035 | 2 | $2.4 \times 10^{-10}$ |
| 37a | *L. pedunculatus* | 166,587,314 | 3 | $3.3 \times 10^{-10}$ |
| 38a | *L. pedunculatus* | 185,499,166 | 6 | $5.9 \times 10^{-10}$ |
| 54a | *M. lupulina* | 258,825,042 | 4 | $2.9 \times 10^{-10}$ |
| 81a | *L. pedunculatus* | 204,485,061 | 2 | $1.8 \times 10^{-10}$ |
| PT | *L. arenarius?* | 213,058,132 | 2 | $1.7 \times 10^{-10}$ |
| **Total:** | | 2,468,379,192 | 43 | $3.2 \times 10^{-10}$ |

We identified 43 high confidence and 24 low confidence putative mutations (see Materials and Methods), and selected 18 high confidence and 8 low confidence mutations for validation with Sanger sequencing. We obtained amplicons of expected lengths for all 18 high confidence mutations, and six low confidence mutations. Sanger sequencing results revealed that all six low confidence mutations were false positives, confirming the suitability of our filtering criteria. All low confidence mutations were assumed to be false positives and excluded from calculation of mutations rate. For high confidence mutations, Sanger sequencing failed for six amplicons despite repeated attempts. From the remaining twelve candidate mutations, we confirmed ten and identified two false-positives (Figure S2 and Table S3). This results in false-positive rate of 16.7%, which is similar to previously reported rates in other insects (Keightley et al. 2014, 2015; Oppold and Pfenninger 2017). With correction for the false positive rate, we estimate the mutation rate per site per haploid genome per generation as 43/(2468379192*2*27)*0.833 = 2.7×10^−10^ (95% CI 1.9×10^−10^-3.5×10^−10^).

Our estimate of the mutation rate in pea aphids is the lowest reported for any insect so far (Keightley et al. 2014, 2015; Yang et al. 2015; Liu et al. 2017; Oppold and Pfenninger 2017). We are still developing an understanding on what drives the differences in mutation rate between species (Bromham 2009), thus we can only speculate as to the mechanisms behind the low mutation rate in pea aphids. One possible explanation is that the low mutation rate is be related to the peculiar life cycle of aphids. Aphid females reproduce by apomictic parthenogenesis (i.e., without meiosis) during most of the year (around 10-15 generations), and with onset of cold conditions a single sexual reproduction event takes place, after which overwintering eggs are laid. Thus, recombination in pea aphids is rare (roughly, 1 meiosis event every 10-15^th^ generation). This may cause increased selection for high fidelity of DNA polymerase in order to alleviate the mutation load resulting from accumulation of deleterious mutations during the parthenogenetic phase. A similar argument has recently been made to explain the low mutation rate observed in giant duckweed *Spirodela polyrhiza* (Xu et al. 2019), a species that reproduces mostly by asexual budding and exhibits the lowest mutation rate of any plant (2.4×10^−10^). On the other hand, water flea *Daphnia pulex* – whose life cycle includes up to 5 apomictic parthenogenetic generations between each sexual reproduction event – exhibits a mutation rate 10x higher than our estimate in pea aphids (Flynn et al. 2017). However, because estimates of mutation rate from other crustaceans are currently missing, it is impossible to judge if this is a low mutation rate compared to other crustaceans. Additional estimates from other species will be needed to test the role of asexual reproduction in mutation rate variation between species.

The size of the pea aphid X-chromosome (around 1/3 of the entire genome (Manicardi et al. 2015; Mandrioli et al. 2017)) provides a rare opportunity to test whether mutations rates in sex chromosomes and autosomes differ. This could potentially explain peculiarities of sex chromosome evolution such as faster differentiation or a preponderant role on speciation (Presgraves 2018). Using available annotation of X-linked region in the pea aphid genome (Jaquiery et al. 2019), we classified de novo mutations as X-linked or autosomal. We found that 11 de novo mutations (out of 42 that could unambiguously be mapped to either X or autosomes, i.e. 26%) occurred on X chromosome (Table S4), a number not significantly smaller than the expected if mutations occur at the same rate in all chromosomes (permutation test, P = 0.2). This suggests similar mutation rates of sex-chromosomes and autosomes, and is consistent with previous results in nematodes (Denver et al. 2012).

### Genomic differentiation suggests more ancient onset of the pea aphid radiation

In order to obtain an estimate of the beginning of the pea aphid radiation, we chose distantly related host races according to the phylogeny in (Fazalova et al. 2018), re-sequenced the genomes of 13 individuals of *L. pratensis* and 12 individuals of *V. cracca* to mean coverage of 20×, and trimmed and mapped sequence reads to the pea aphid genome (on average 88.3% of reads were mapped to the reference genome, Table S1). To alleviate the effects of selection along the genome, and sex-chromosome specific biases, we extracted only 4-fold degenerate sites from autosomal genes for analysis (2,116,329 sites). Similar levels of genome-wide synonymous polymorphism were found in both host races: θ = 0.0042 and 0.0043, for *L. pratensis* and *V. cracca*, respectively. Average genomic divergence between the two host races was estimated as *d*_*XY*_ = 0.0086, and average differentiation along the genome as *d*_*a*_ = *d*_*XY*_ − (*θ*_*X*_ + *θ*_*Y*_)/2 = 0.0086 − (0.0043 + 0.0042)/2 ≈ 0.0044. We can use this simple formula which accounts for ancestral polymorphism because gene flow is very unlikely between *L. pratensis* and *V. cracca*, with evidence of strong intrinsic reproductive isolation (Fazalova et al. 2018) and no hybrids found in the wild (Peccoud et al. 2009a).

Using our estimates of the spontaneous mutation rate in pea aphid and genetic divergence between *L pratensis* and *V.* cracca, we calculated the age of divergence of the pea aphid complex as 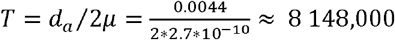 generations (95%CI 6 286,000 − 11 579,000). To convert the number of the generations into years, we assumed 15 generations per year: this results into a time of split of about 543,000 years (95% CI 419,000– 772,000). This estimate is conservative, as it includes the maximal number of asexual generations per year suggested for aphids: 14 (Loxdale and Balog 2018). We added one more generation to this estimate to calculate the total number of generations per year, as we assume that sexual generation is likely to have similar mutation rate. Even if the mutation rate is higher in sexual generations, it does not affect our estimate of the divergence time very strongly, as aphids have a single sexual generation per year, and the rest of the life cycle consists of asexual generations. If we assume that the pea aphid mutation rate during sexual generation is similar to other insects (i.e. 3e-9, which is an average of the estimates obtained from (Keightley et al. 2014, 2015; Yang et al. 2015; Liu et al. 2017; Oppold and Pfenninger 2017)), we can estimate the mutation rate which accounts for contribution of sexual reproduction every 15^th^ generation of the life cycle as 2.27e-10*(14/15) + 3e-9*(1/15) = 4.12e-10, and the age of the pea aphid divergence would be about 356,000 years.

Our estimate of the age of the pea aphid radiation contrasts with previous report of very recent and rapid diversification (Peccoud et al. 2009b), which was based on the mutation rate of maternally transmitted endosymbiont *Buchnera aphidicola* (Moran et al. 2009). However, *B. aphidicola* genome behaves as a single gene, without recombination (Shigenobu et al. 2000) and exclusive maternal inheritance (Tóth 1933). Reconstruction of evolutionary histories and estimation of divergence times from such genes have been shown to be unreliable and strongly affected by processes such as selective sweeps or drift (Shaw 2002; Ballard and Whitlock 2004). Given our estimate is based on over 2 million sites sampled genome-wide, we expect our new estimate of the divergence age of the pea aphid to be much more reliable than previous estimates.

Our results contradict the Post-Pleistocene timing of speciation in the pea aphid complex and rule out the effect of anthropogenic Neolithic agriculture on their diversification. The older age of the radiation 543,000 years (95% CI 419,000– 772,000) raises the possibility that pea aphids experienced numerous Pleistocene habitat fragmentations (Hewitt 2004). They might have been trapped in separate refugia (Stewart et al. 2010) on novel host plants, and undergone adaptation and divergence without the impeding effect of gene flow. This implies that patterns of gene flow present in pea aphids can be more parsimoniously explained by allopatric isolation and secondary contact than divergence with gene flow. This has been suggested before by some authors (e.g., Futuyama 2008; Bierne et al. 2013; Harrison and Larson 2016) but remains largely unappreciated in the large body of work on pea aphid speciation.

Our results shed new light into the diversification of the pea aphid complex, and call for reassessment of current understanding of this study system. It will be especially important to understand how much geographic separation has contributed to the divergence of the closely related pea aphid host races (specialized on *Medicago sativa* and *Trifolium pratense*). Thus far, patterns of genomic differentiation between pea aphid host races have been interpreted in light of a scenario of speciation with gene flow. In particular, genome scans studies have identified highly differentiated genomic regions between host races and interpreted those as regions involved in adaptation to host plants, because under speciation with gene flow genes responsible for adaptation are expected to show elevated differentiation. Furthermore, the identification of large genomic regions of high divergence around quantitative trait loci in pea aphids has inspired the development of the theory of divergence hitchhiking, that is the process through which initial selection on only a few loci can extend divergence to larger genomic regions (Via and West 2008). However, under a scenario of allopatric isolation, any genomic region has the potential to differentiate without necessarily being involved in reproductive isolation or adaptation (e.g., Feder et al. 2013). Thus, both genome scans results and divergence hitchhiking models need to be reassessed in light of the potential role of geographic isolation in divergence of pea aphid host races. Finally, our results raise warning for other study systems as well – especially for those which are assumed to have recently divergence with ongoing gene flow. Only lineage-specific estimates of mutation rates, together with genome-wide estimates of divergence – as performed here – will allow accurate inference of evolutionary histories and a more complete understanding of the processes driving diversification in these systems.

## Materials and Methods

### Collection and rearing of aphids

We sampled overlapping populations of pea aphids from several wild Fabaceae host plants in summers of 2015-2016 around Oxford, UK (Table S5). We genotyped all aphids collected with 14 microsatellite loci (Peccoud et al. 2009a) to confirm the host race assignment (except the pea aphid line collected from *Lotus arenarius*), and reared them as parthenogenetic lines in the lab at 14±1°C with a 16L: 8D photoperiod, on leaves of *Vicia faba* (Sutton variety, replaced weekly) placed in 1% agarose gel in Petri dishes.

### Mutation accumulation experiment

For the mutation accumulation experiment, in May 2017 we established twelve parthenogenetic lines: three from *Lathyrus pratensis*, three from *Lotus corniculatus*, three from *Lotus pedunculatus*, two from *Medicago lupulina* and one from *Lotus arenarius*. To establish each line, before starting the mutation accumulation experiment we isolated a single asexual female and waited until reproduction, at which point we randomly selected six offspring to establish the first generation and used the remaining offspring for DNA extraction. Each generation thereafter was established by randomly selecting six offspring, ensuring low population sizes throughout the experiment. We continued these transfers until July-September 2018, at which point all lines had gone through 28 generations. For the penultimate generation, we isolated a random single adult female and collected its offspring for DNA extraction.

### DNA extraction and sequencing

For each aphid line and generation, we extracted DNA from pools of six individuals (sisters) to obtain sufficient quantity for preparation of DNA libraries for whole-genome resequencing. Aphids were frozen in −80C, processed with TissueLyser (Qiagen), and DNA extracted following the Supplementary Qiagen protocol (Purification of total DNA from insects using the DNeasy® Blood & Tissue Kit). We treated DNA extractions with 2 μl of RNAase A (ThermoScientific) and purified them with Monarch PCR & DNA Cleanup Kit (New England Biolabs). We assessed DNA quantity and quality with Qubit 3.0 (Thermofisher), Nanodrop (Thermofisher) and gel electrophoresis (0.7% agarose gel, TAE buffer). Preparation and sequencing of DNA libraries were performed by Novogene (Hong Kong). For the mutation accumulation lines we used TruSeq libraries (Illumina), and sequenced with 150 bp paired-end reads to > 30× coverage per line and generation. For the analysis of genetic differentiation, we used NEBNext Ultra library kits (New England Biolabs) and sequenced to >15× coverage per line.

### Identification of candidate mutations, validation, estimation of mutation rate

We quality-controlled raw sequencing data with fastqc (Babraham Bioinformatics) and trimmed sequencing adaptors and low quality reads with trim-galore (https://www.bioinformatics.babraham.ac.uk/projects/trim_galore/) using Cutadapt (Martin 2011): we discarded bases with quality below 20 and discarded read pairs if any was shorter than 36 bp after trimming. We mapped the trimmed reads to the reference genome version Acyr_2.0 using bwa (Li and Durbin 2009), with default parameter values and the -M option (which marks shorter split hits as secondary), and removed duplicate reads with samtools rmdup (Li et al. 2009). We realigned regions around indels using the GATK (REF) indel realignment tool. At each step of data processing, summary files were inspected in multiqc (Ewels et al. 2016).

We performed variant calling with samtools and bcftools (Li et al. 2009) with the multi-allele caller (-m option) and discarding reads with mapping quality below 20 and bases with base quality below 20. We filtered variants with bcftools, excluding SNPs with low quality (<20) and with too low (<20) or too high (> 2.5 times average depth for each sample) depth. We further filtered heterozygous SNPs with fewer than 2 reads supporting each allele, SNPs near indels (< 3 bp), indels separated by 10 or fewer base pairs and SNPs supported by fewer than 2 reads in each direction (for heterozygous genotype calls that include the reference allele, i.e. GT=0/1). Versions of the software are provided in Table S6.

In order to obtain a first set of candidate *de novo* mutations in each line, we identified SNPs covered by at least 20 reads in each generation and with different genotype calls in generations 0 and 28 (start and end of mutation accumulation lines) considering only base substitutions (i.e. ignoring indels). Candidate mutations were further filtered out if reads supporting both alleles are present in both lines (generations 0 and 28) and if genotypes of the two samples imply more two different alleles. These steps were implemented in custom scripts (available from the authors) that use the mpileup files produced by samtools for generations 0 and 28.

We further filtered this set of candidate mutations, by manually inspecting the alignment files in IGV (Thorvaldsdottir et al. 2013) for the following artefacts suggesting read mapping errors:

1. Candidate mutations do not show consistent linkage to other polymorphic sites on the same read (Long et al. 2016), as expected due to apomictic parthenogenesis.
2. Reads supporting candidate mutations appear both in Generation 1 and Generation 28 (for cases where read, base or mapping quality were too low and thus not considered in previous steps).
3. Reads supporting putative mutations have many substitutions, which are false positives according to the filtering criterion 2.
4. SNPs supported by fewer than two reads in each direction (for heterozygotes genotype calls that do not include the reference allele, i.e. GT=1/2).
5. Candidate mutation can be resolved by indel realignment (Figure S3 in Keightley et al. 2014).

With these filters we obtained two sets of candidate *de novo* mutations. High confidence mutations passed all filters and were expected to be true *de novo* mutations. Low confidence mutations passed the filtering criteria, but were considered unlikely either because (1) reads supporting the candidate mutation were clipped or had many indels/substitutions (Figure S1 in Keightley et al. 2014), (2) more than one candidate mutation occurred within 50 bp region, or (3) candidate mutation implied a change from heterozygous to homozygous state. We also included in the low confidence set one candidate mutation for which a single read supporting the new allele was also found in the Generation 1, to test if the filtering criterion 2 was too stringent. Candidate mutations after each step of filtering are shown in Table S7.

For all candidate mutations (low and high confidence) we retrieved FASTA files (based on the reference genome Acyr. 2.0) with 600 bp flanking the candidate mutations and attempted to design primers for these regions, using Primer3Plus (Untergasser et al. 2007). Primers were excluded if they would map to other scaffolds of the genome, as inferred with Primer-BLAST (Ye et al. 2012). This resulted in 25 primer pairs, 18 targeting regions around high confidence mutations, and 7 around low confidence mutations, which were used for PCR amplification of both generations (Table S8). PEG clean-up and Sanger sequencing of the PCR product (BigDye 3.1, Applied Biosystems capillary 3730XL DNA Analyzer) were performed by the Sequencing Core of the Department of Zoology, University of Oxford.

We estimated the mutation rate per haploid genome using the number of high confidence mutations and the number of callable sites (total number of sites with adequate depth of sequencing in both generation 0 and generation 28, as explained above) for each line. In addition, we estimated mutation rate for both X and Autosomes, using the assignment from (Jaquiery et al. 2019).

### Estimation of time of divergence

For the analysis of genomic differentiation, we chose distantly related host races according to the phylogeny in (Fazalova et al. 2018), and included 13 individuals of *L. pratensis* and 12 individuals of *V. cracca*. Quality control, trimming, removal of duplicate reads, mapping, indel realignment and variant calling were performed in the same way as for the mutation accumulation analysis (see above). Filtering was less stringent compared to the mutation accumulation pipeline due to lower coverage: obtained SNPs were filtered to remove low quality SNP calls (< 15); SNPs with low depth (< 8 reads); heterozygous SNPs with less than 2 reads supporting each allele; SNPs near indels (< 3 bp); indels separated by 10 or fewer base pairs; and SNPs with high depth (more than 2.5x the average depth of coverage of each sample). We used the option to report homozygous-reference blocks for each individual with a minimum depth of 8 reads. Versions of the software are provided in the Table S6.

In order to obtain an estimate of divergence that is least affected by selection, for analysis of divergence we used only 4-fold degenerate sites from each gene. We used vcf2fas (available from https://github.com/brunonevado/vcf2fas) to obtain FASTA files for each scaffold, and extracted coding sequences (CDS) according to the available annotation of the pea aphid genome obtained from GenBank (GCF_000142985.2_Acyr_2.0_genomic.gff). We assigned CDS to X-chromosomes and Autosomes according to (Jaquiery et al. 2019). For each gene, we extracted and concatenated 4-fold degenerate sites from all exons using custom scripts (available from the authors).

In order to calculate genetic differentiation between *L. pratensis* and *V. cracca*, while taking into account ancestral polymorphism, we calculated population polymorphism (Watterson theta (Watterson 1975)) and absolute divergence (*d*_*XY*_, (Nei and Li 1979)) using Popgenome (Pfeifer et al. 2014), and used the formula of Nei and Li (Nei and Li 1979) *d*_*a*_ = *d*_*XY*_ − (*θ*_*X*_ + *θ*_*Y*_)/2, where *d*_*XY*_ is absolute divergence between two populations, and *θ*_*X*_ and *θ*_*Y*_ is polymorphism within *L. pratensis* and *V. cracca* respectively. To convert our estimate of genetic differentiation into absolute age of divergence (in generation) we used then used the formula *T* = *d*_*a*_/2*µ*, where *µ* is the mutation rate (per site per generation).

## Author Contributions

VF designed the study, collected the data and performed the mutation accumulation experiment. VF and BN performed the analysis. VF wrote the manuscript with input from BN. Both authors agree with the content of the manuscript.

## Acknowledgments

We are grateful to Charles Godfray for useful comments on the manuscript. VF is grateful to Ciara Mann for assistance in collecting and maintaining aphids, to Jason Hogg for maintenance of controlled temperature rooms and to the department of Plant Sciences of the University of Oxford for hosting after the closure of the Tinbergen building. The authors acknowledge the use of the University of Oxford Advanced Research Computing (ARC) facility in carrying out this work. http://dx.doi.org/10.5281/zenodo.22558. This project has received funding from the European Union’s Horizon 2020 research and innovation programme under the Marie Skłodowska-Curie grant agreement no. 659847 (to VF), and from the John Fell Fund grant no. 0005545 from the University of Oxford (to VF).

