## Supplemental Files for "Low spontaneous mutation rate and Pleistocene radiation of pea aphids"

**Table S1. Summary of sequencing data for each sample**

| Host_race | Sample | Raw_reads each_end (x10$^6$) | Coverage | Percentage_of reads_mapped |
|---|---|---|---|---|
| *Lat. pratensis* | A16_19a | 35.3 | 22.82 | 93.11 |
| *Lat. pratensis* | A16_26a | 26.8 | 17.33 | 93.79 |
| *Lat. pratensis* | A16_62a | 40.5 | 26.19 | 92.6 |
| *Lat. pratensis* | A16_80a | 32.3 | 20.88 | 93.42 |
| *Lat. pratensis* | A16_107a | 27.7 | 17.91 | 92.02 |
| *Lat. pratensis* | A16_110a | 40.4 | 26.12 | 93 |
| *Lat. pratensis* | A16_116a | 28.6 | 18.49 | 93.65 |
| *Lat. pratensis* | A16_120a | 31.1 | 20.11 | 93.33 |
| *Lat. pratensis* | A16_138 | 28.6 | 18.49 | 92.96 |
| *Lat. pratensis* | A16_161b | 28.1 | 18.17 | 93.69 |
| *Lat. pratensis* | A29d | 34.3 | 22.18 | 94.73 |
| *Lat. pratensis* | A61b | 28.1 | 18.17 | 95.18 |
| *Lat. pratensis* | A70a | 34.3 | 22.18 | 93.62 |
| *V. cracca* | A16_28a | 32.4 | 20.95 | 83.48 |
| *V. cracca* | A16_61a | 28.6 | 18.49 | 87.53 |
| *V. cracca* | A16_63a | 28.9 | 18.69 | 83.19 |
| *V. cracca* | A16_84a | 38.3 | 24.76 | 86.29 |
| *V. cracca* | A16_95a | 29.6 | 19.14 | 88.9 |
| *V. cracca* | A16_105a | 38 | 24.57 | 79.48 |
| *V. cracca* | A16_131a | 31.8 | 20.56 | 85.32 |
| *V. cracca* | A642 | 27.9 | 18.04 | 73.62 |
| *V. cracca* | A56b | 34.1 | 22.05 | 79.25 |
| *V. cracca* | A60c | 30.5 | 19.72 | 83.55 |
| *V. cracca* | A65a | 29.1 | 18.81 | 82.08 |
| *V. cracca* | A67a | 29.8 | 19.27 | 80.8 |
| | | **Average:** 31.8 | **Average:** 20.56 | **Average:** 88.34 |
| *M. lupulina* | B05b_1 | 55.5 | 35.88 | 88.17 |
| *Lot. corniculatus* | B09b_1 | 55.1 | 35.63 | 92.18 |
| *Lat. pratensis* | B107a_1 | 55.5 | 35.88 | 92.46 |
| *Lot. corniculatus* | B118a_1 | 63.4 | 40.99 | 93.27 |
| *Lat. pratensis* | B120a_1 | 55.4 | 35.82 | 91.34 |
| *Lot. corniculatus* | B122a_1 | 67.7 | 43.77 | 91.89 |
| *Lat. pratensis* | B29d_1 | 57.2 | 36.98 | 92.39 |
| *Lot. pedunculatus* | B37a_1 | 61.6 | 39.83 | 90.16 |
| *Lot. pedunculatus* | B38a_1 | 51.4 | 33.23 | 89.48 |

**Table S1 (continued)**

| Host_race | Sample | Raw_reads each_end (x10$^6$) | Coverage | Percentage_of reads_mapped |
|---|---|---|---|---|
| *M. lupulina* | B54a_1 | 71 | 45.91 | 88.12 |
| *Lot. peduncualtus* | B81a_1 | 56.5 | 36.53 | 89.64 |
| *Lot. arenaruis?* | BPT_1 | 68.5 | 44.29 | 88.61 |
| *M. lupulina* | E05b_28 | 60.1 | 38.86 | 83.00 |
| *Lot. corniculatus* | E09b_28 | 53.5 | 34.59 | 93.38 |
| *Lat. pratensis* | E107a_28 | 62.9 | 40.67 | 94.16 |
| *Lot. corniculatus* | E118a_28 | 58.9 | 38.08 | 95.70 |
| *Lat. pratensis* | E120a_28 | 66.5 | 43.00 | 93.68 |
| *Lot. corniculatus* | E122a_28 | 56.4 | 36.47 | 90.83 |
| *Lat. pratensis* | E29d_28 | 52.3 | 33.81 | 94.03 |
| *Lot. pedunculatus* | E37a_28 | 50.5 | 32.65 | 90.57 |
| *Lot. pedunculatus* | E38a_28 | 63.6 | 41.12 | 92.07 |
| *M. lupulina* | E54a_28 | 58.6 | 37.89 | 87.54 |
| *Lot. peduncualtus* | E81a_28 | 58 | 37.50 | 90.20 |
| *Lot. arenaruis?* | EPT_28 | 51.8 | 33.49 | 91.73 |
| | | **Average:** 58.83 | **Average:** 38.04 | **Average**: 91.02 |

**Figure S2. Validation of candidate mutations**

Line: 05b; Scaffold59:515379; A->A/G – High confidence – Confirmed

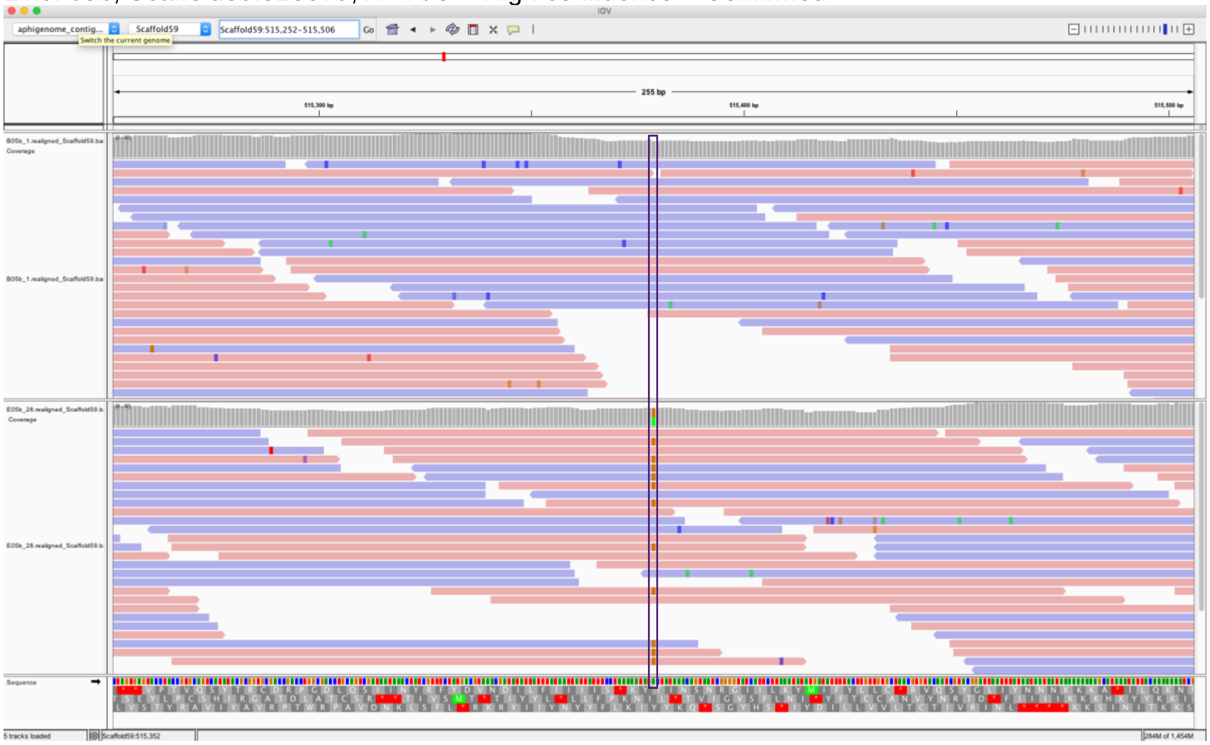


Generation 1:

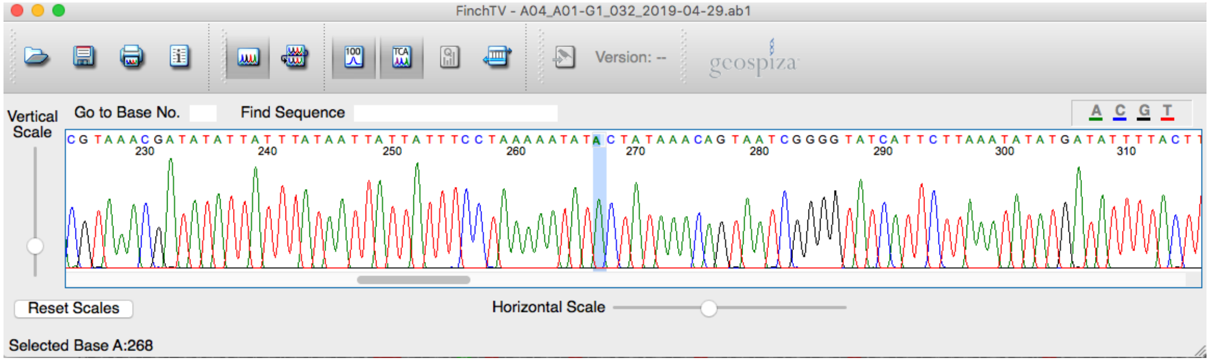


Generation 28:

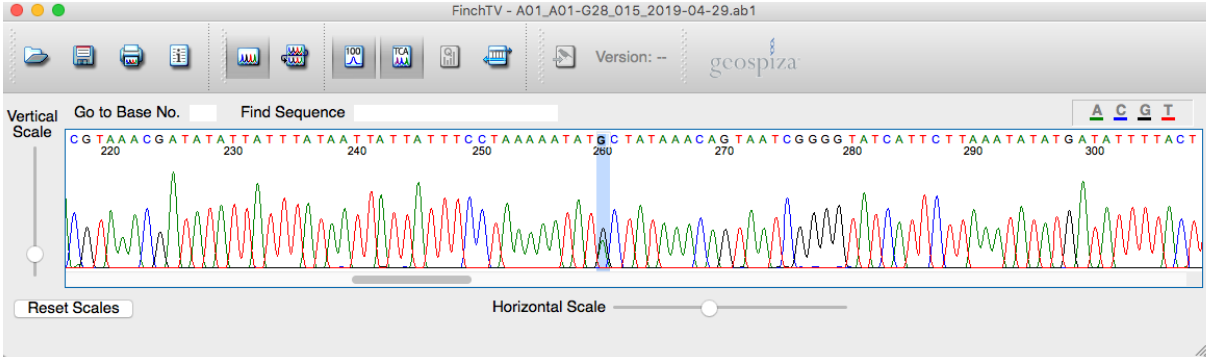

**Figure S2 (continued).** Line 09b; Scaffold153:382214; T->T/G; High confidence – Not Confirmed

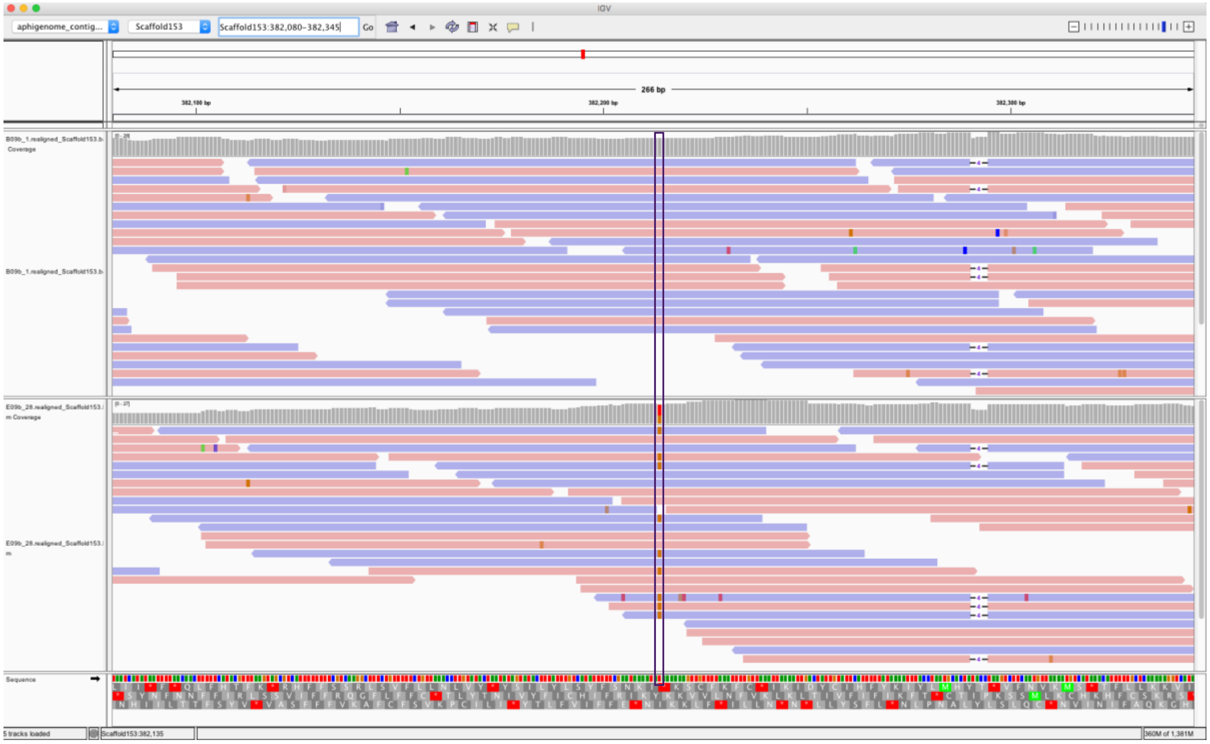


Generation 1:

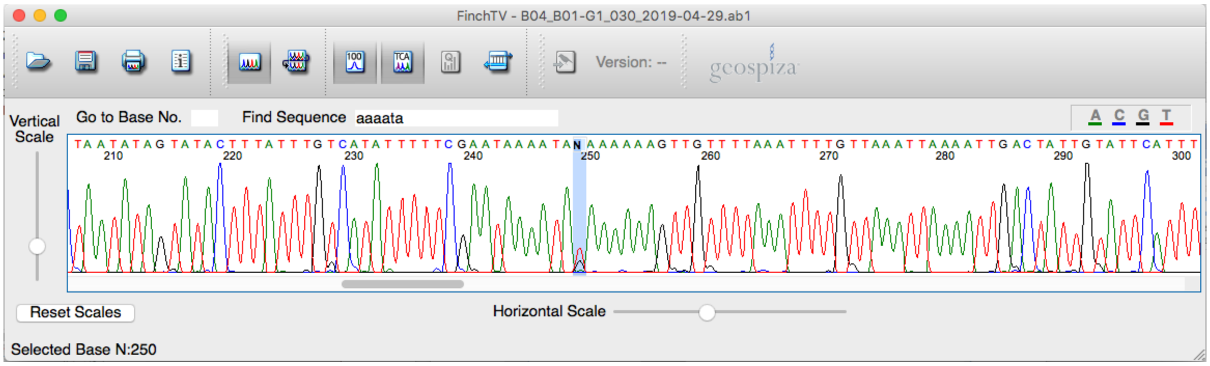


Generation 1 (second Sanger sequencing attempt):

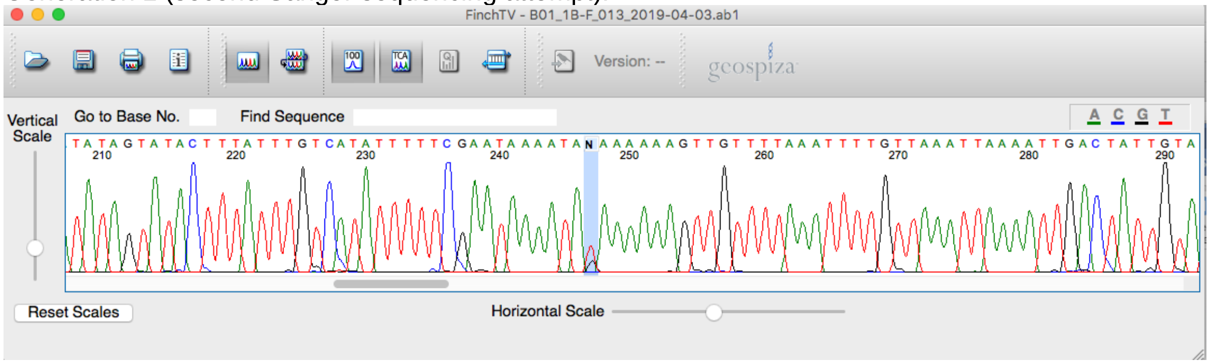

**Figure S2 (continued).** Line: 09b; Scaffold41:243055; T->T/A – High confidence – Confirmed

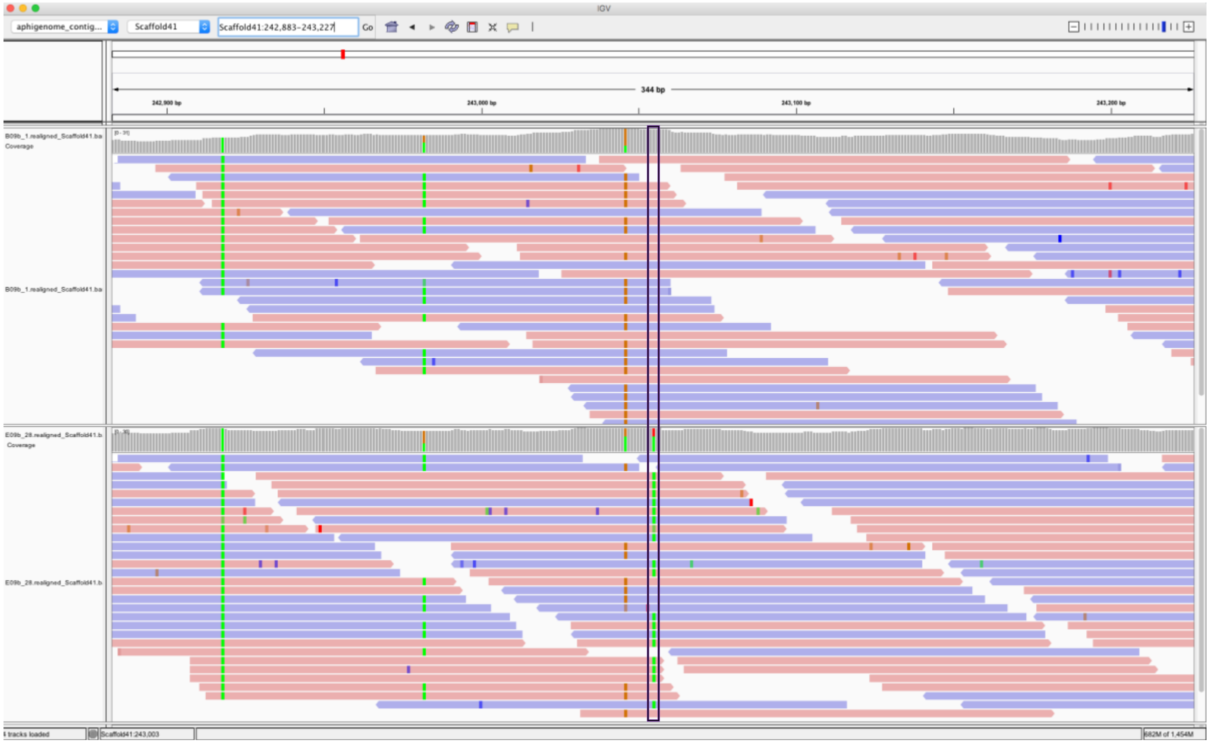


Generation 1:

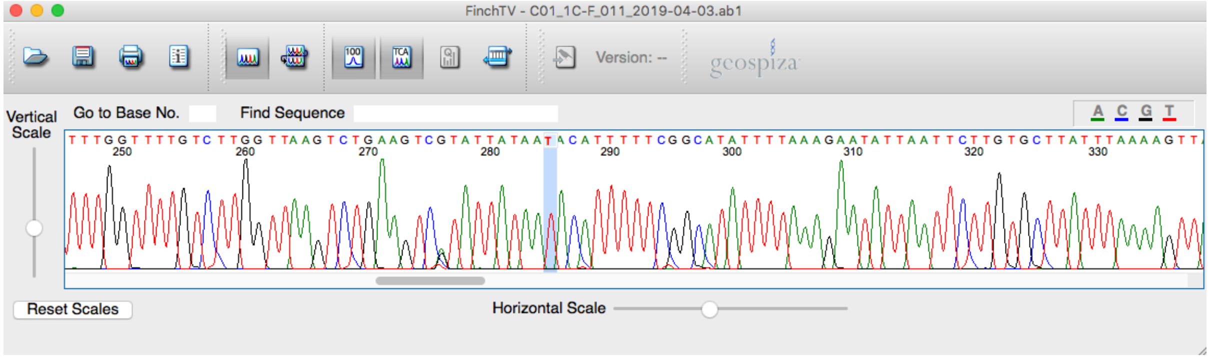


Generation 28:

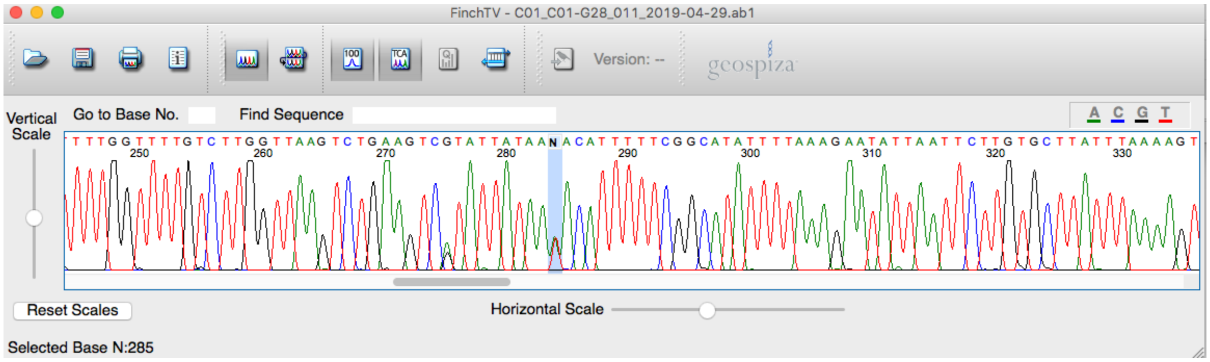

**Figure S2 (continued).** Line 120a; Scaffold1559:38127; C->C/T – High confidence – Confirmed

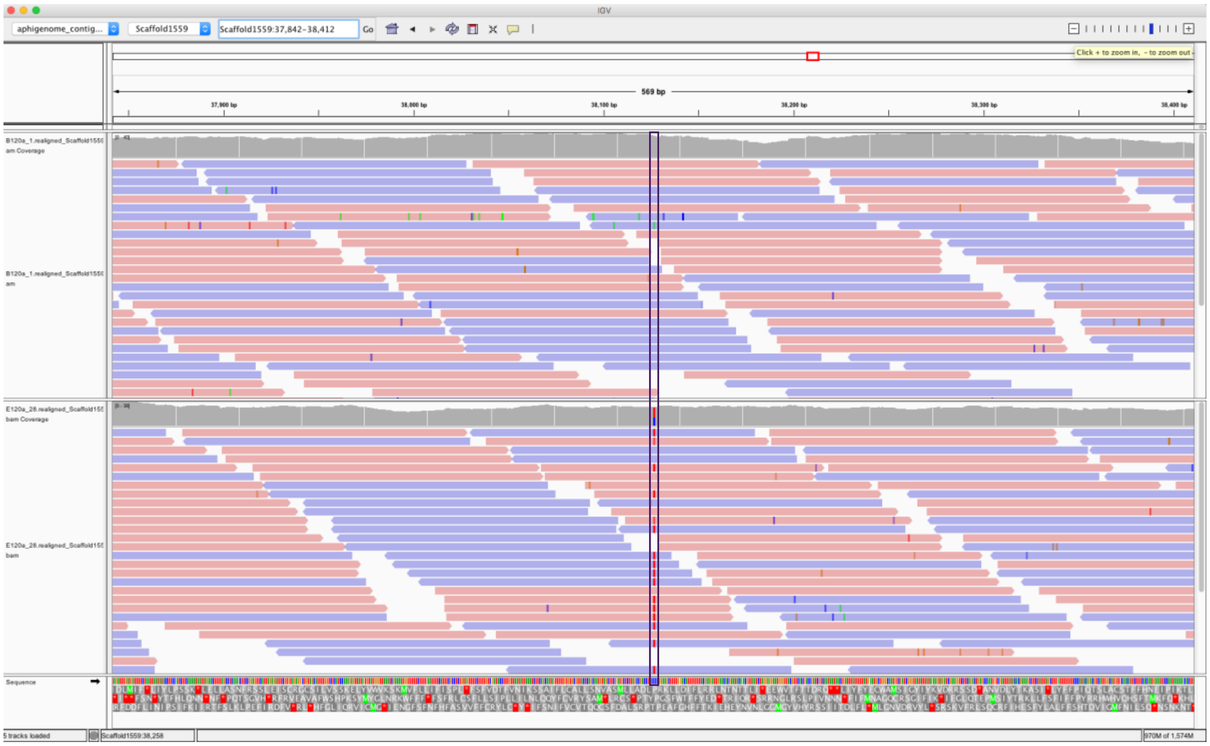

Generation 1:

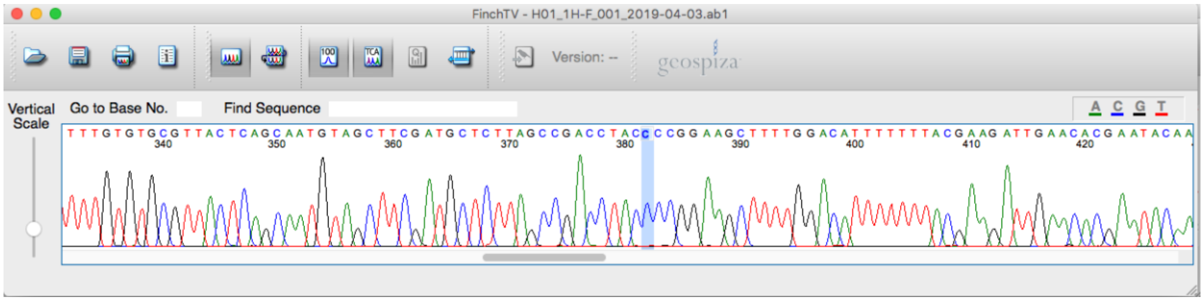


Generation 28:

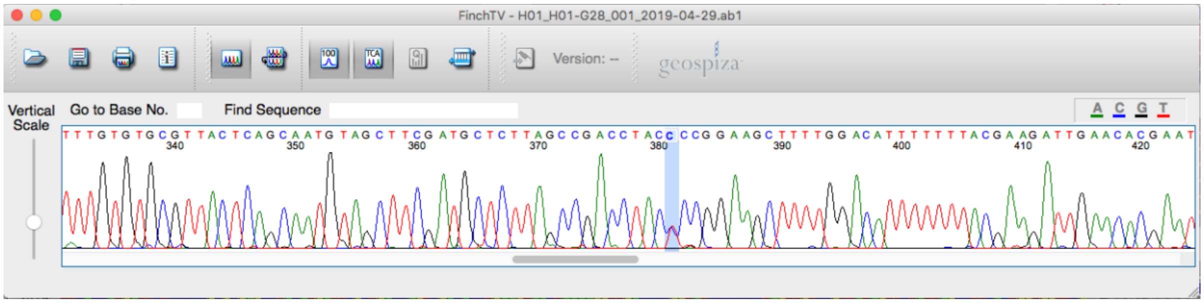

**Figure S2 (continued).** Line: 122a; Scaffold25:659741; G->G/A – High confidence – Confirmed

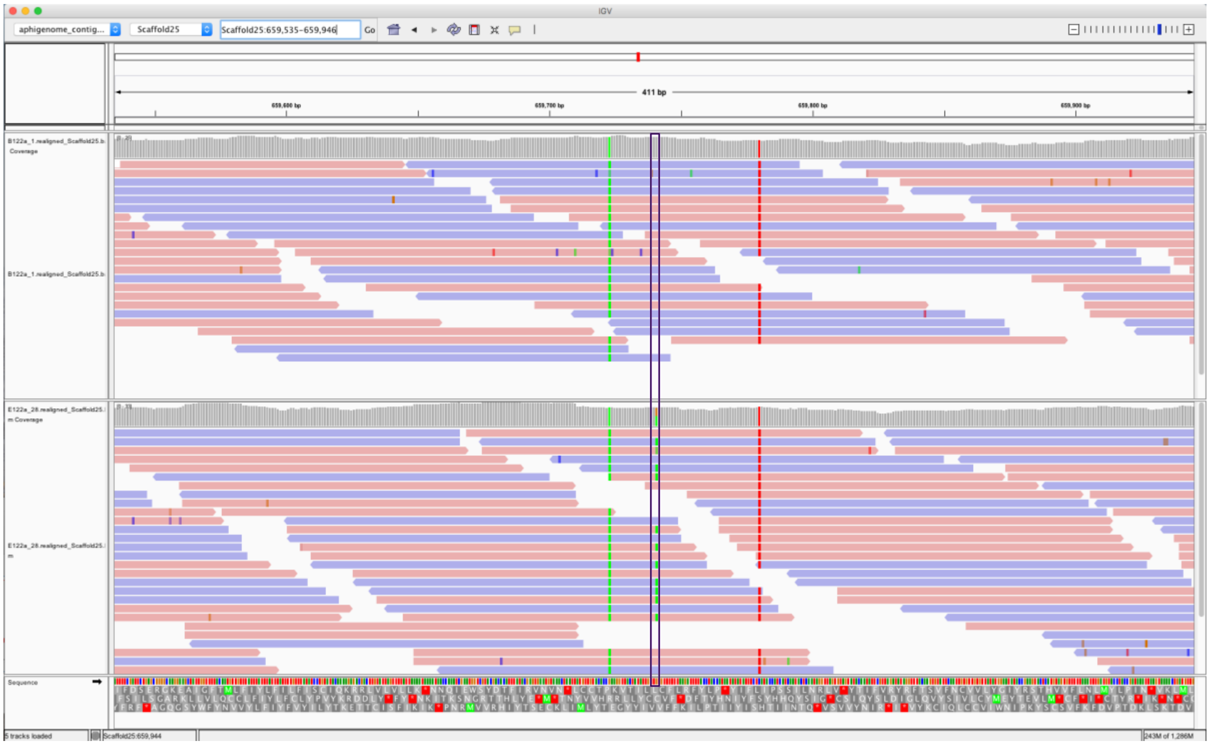


Generation 1:

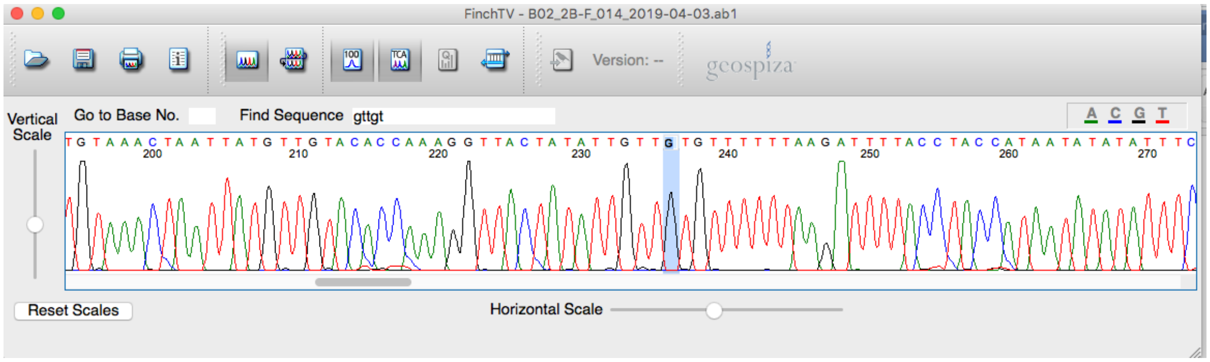


Generation 28:

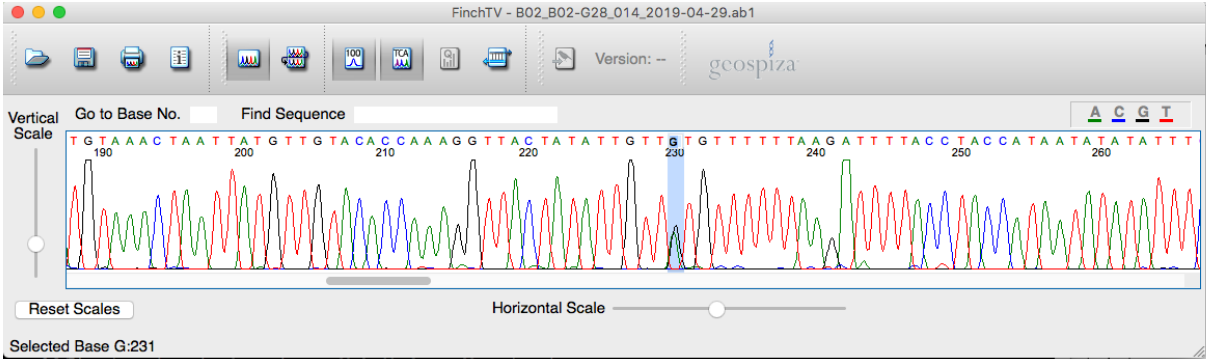

**Figure S2 (continued).** Line: 29d; Scaffold20:513622; C->C/T – High confidence – Confirmed

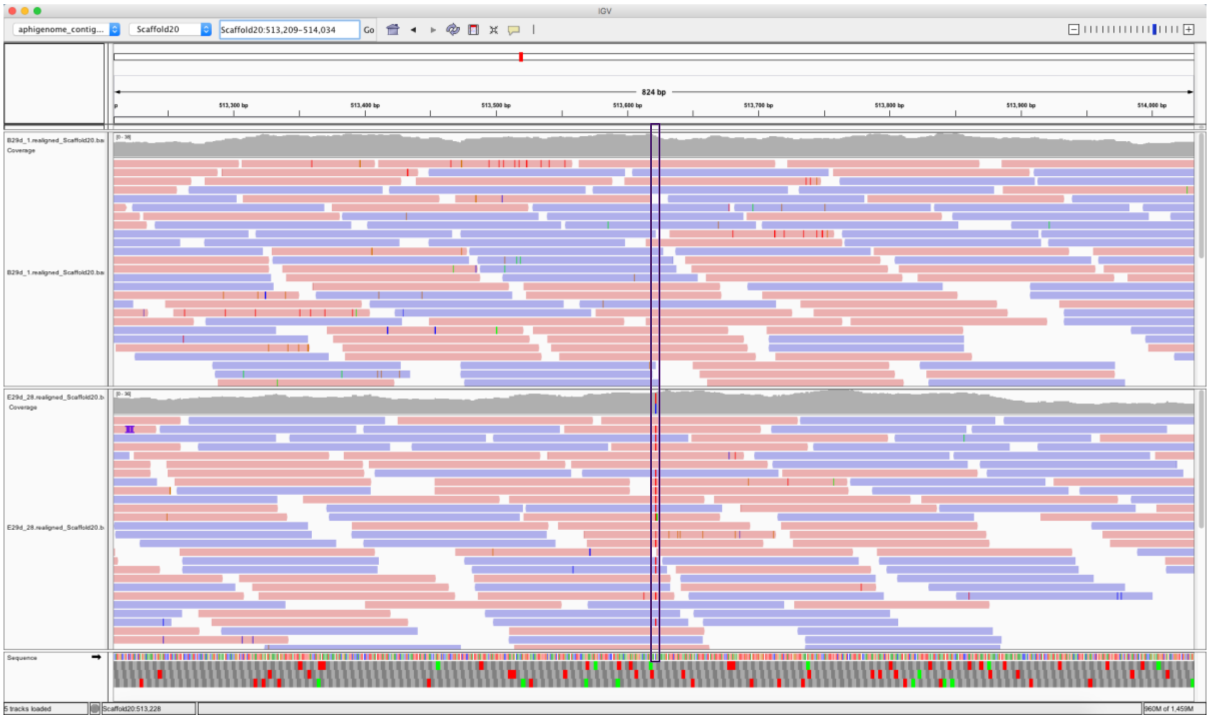


Generation 1:

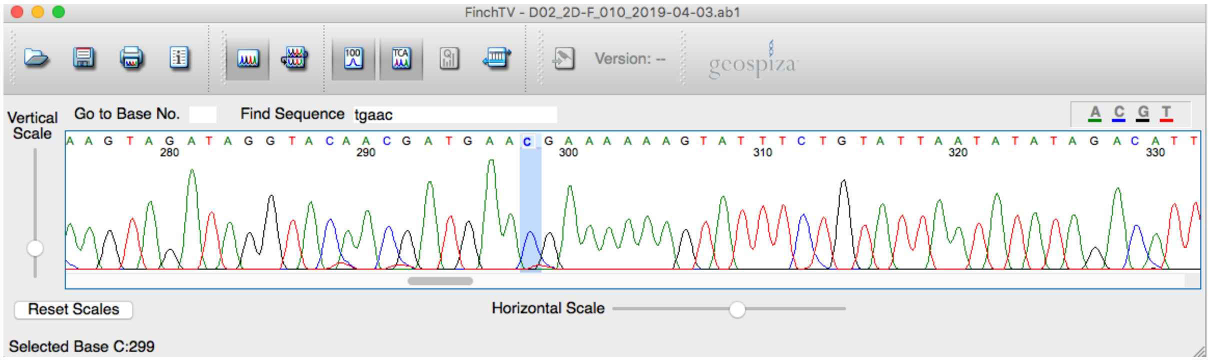


Generation 28:

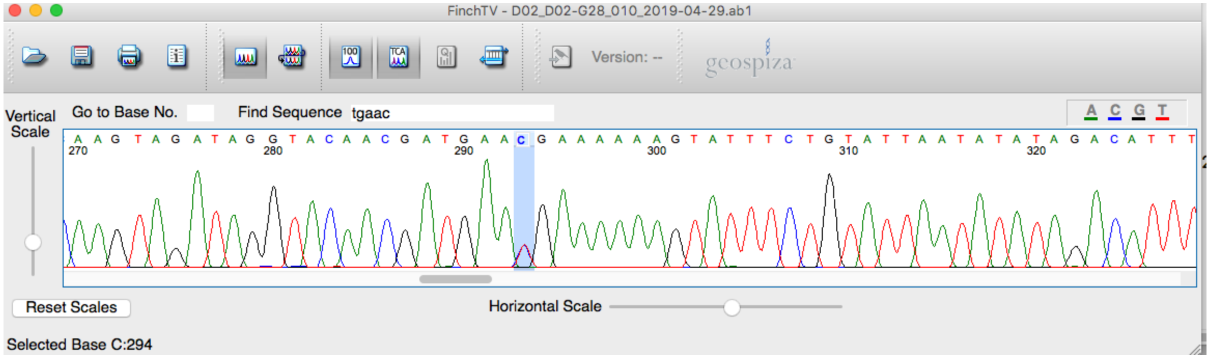

**Figure S2 (continued).** Line 37a; Scaffold68:678872; G->G/A – High confidence – Confirmed

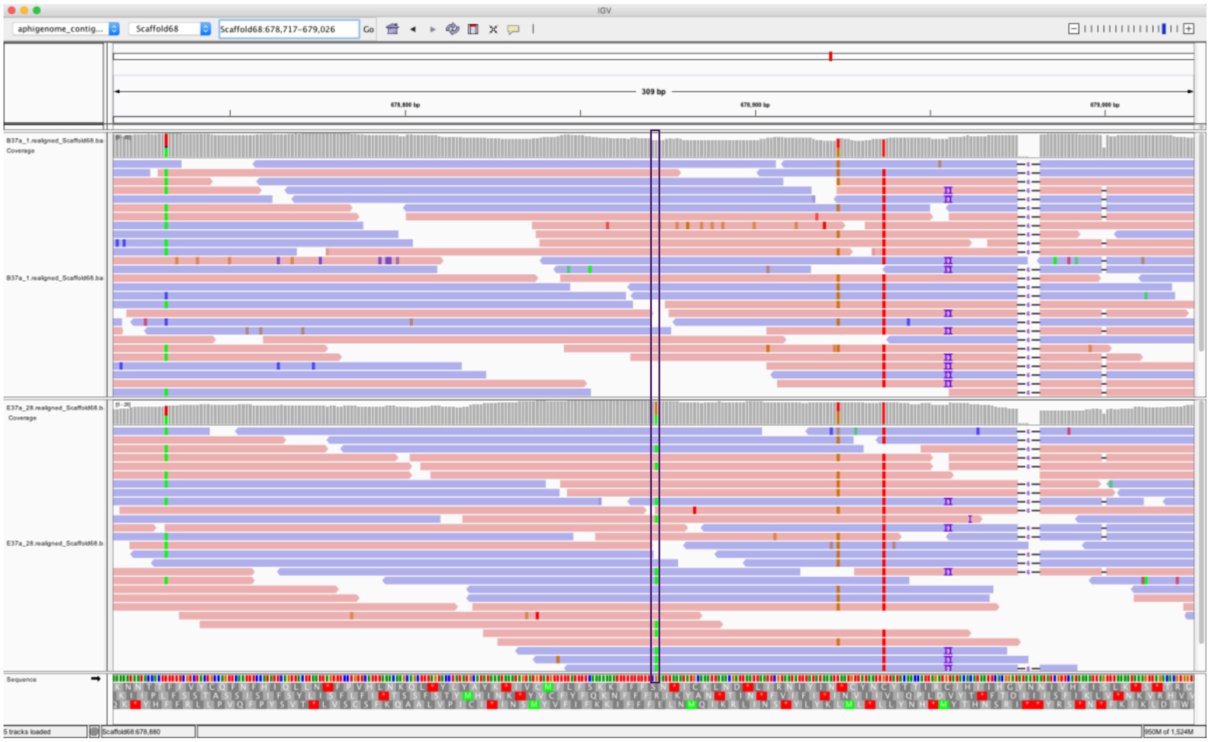


Generation 1:

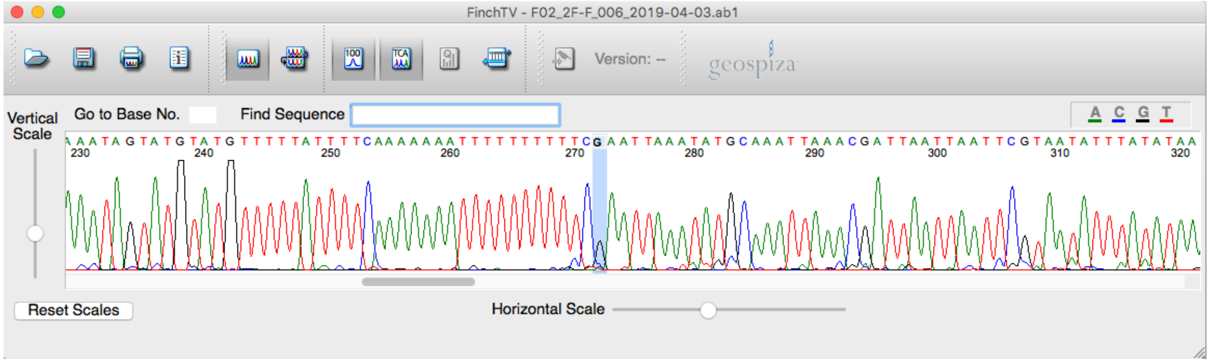


Generation 28:

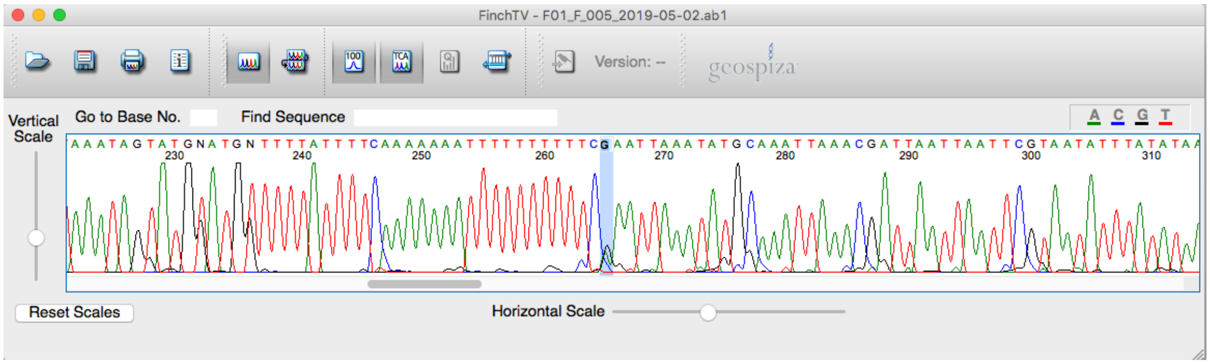

**Figure S2 (continued).** Line 38a; Scaffold359:316424; C->C/A – High confidence – Confirmed

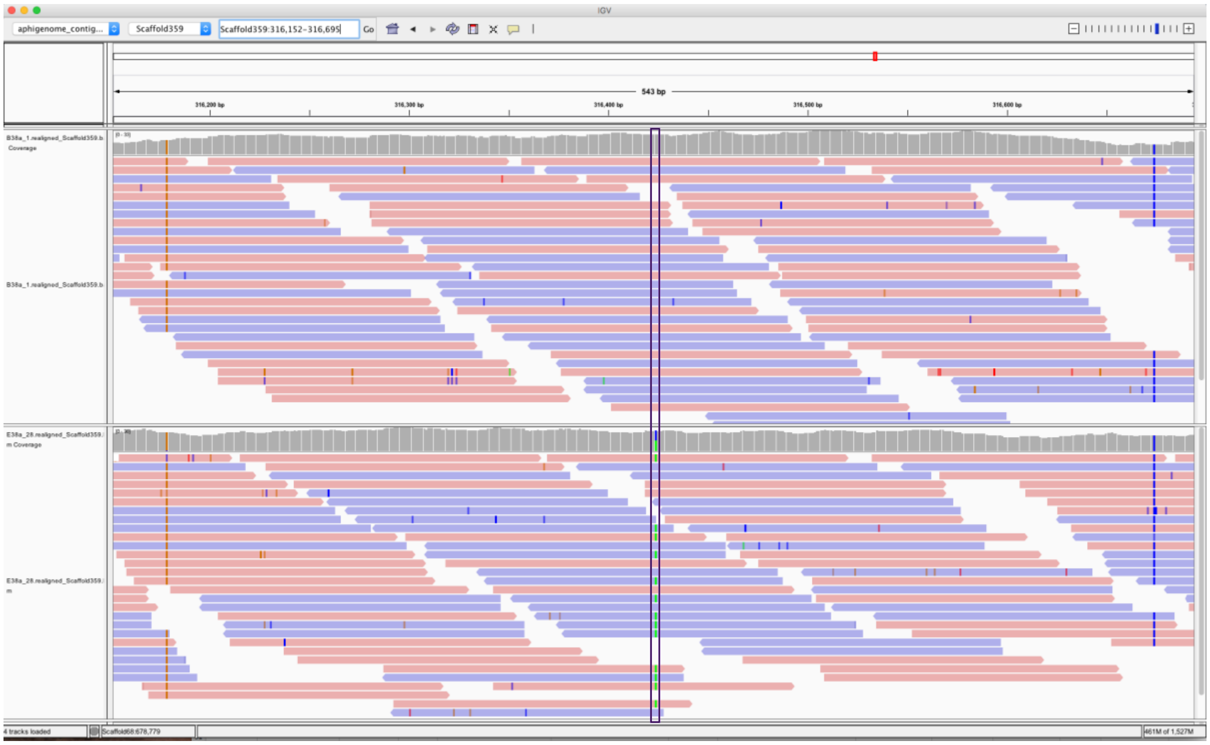


Generation 1:

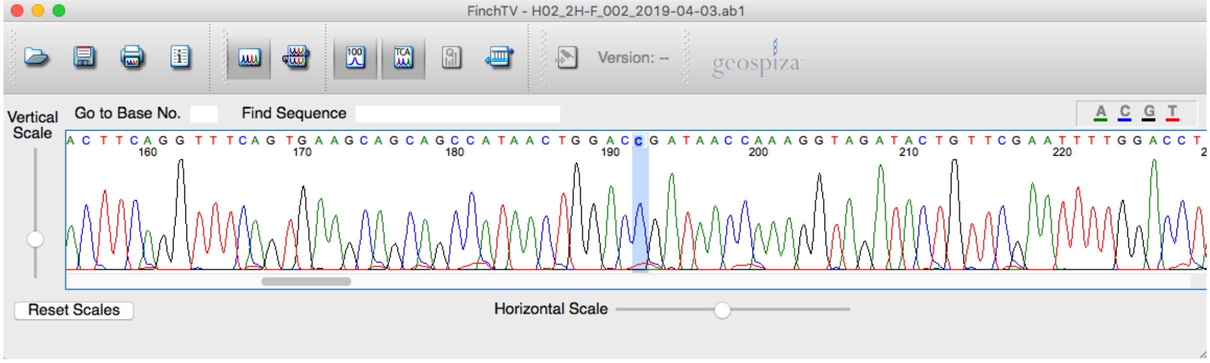


Generation 28:

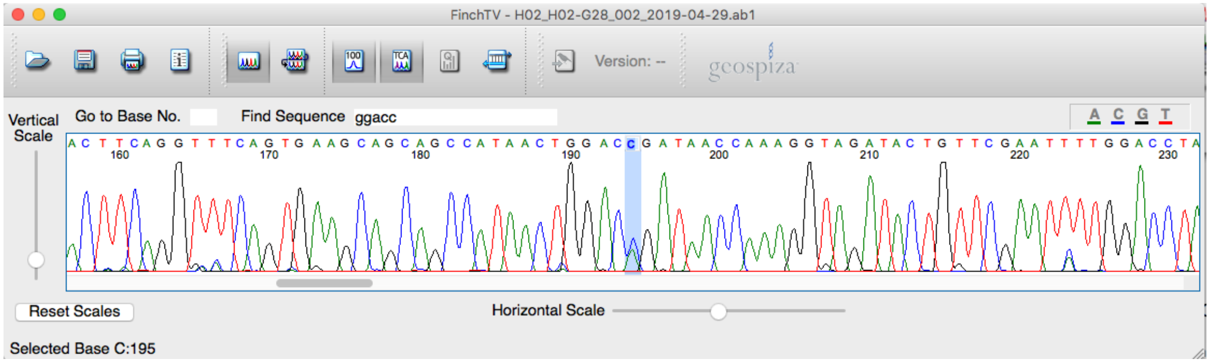

**Figure S2 (continued)**. Line 38a; Scaffold42:194414; C->C/T– High confidence – Not Confirmed

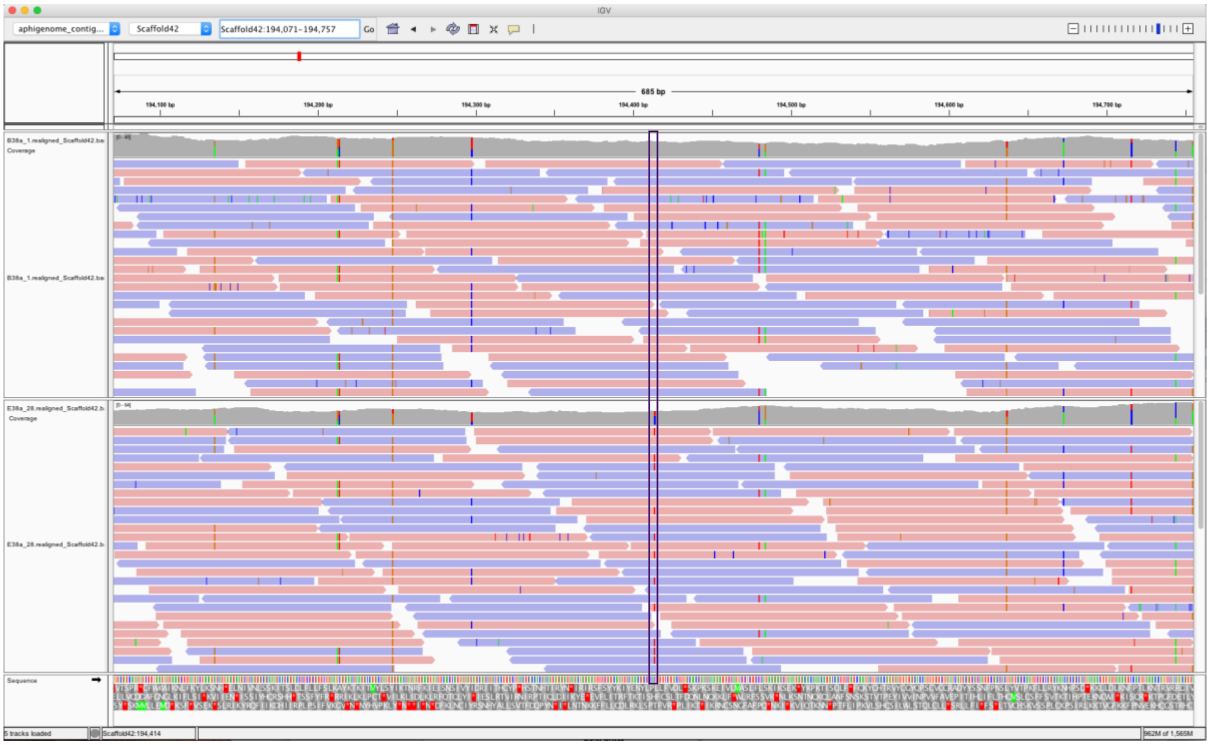


Generation 1(forward primer):

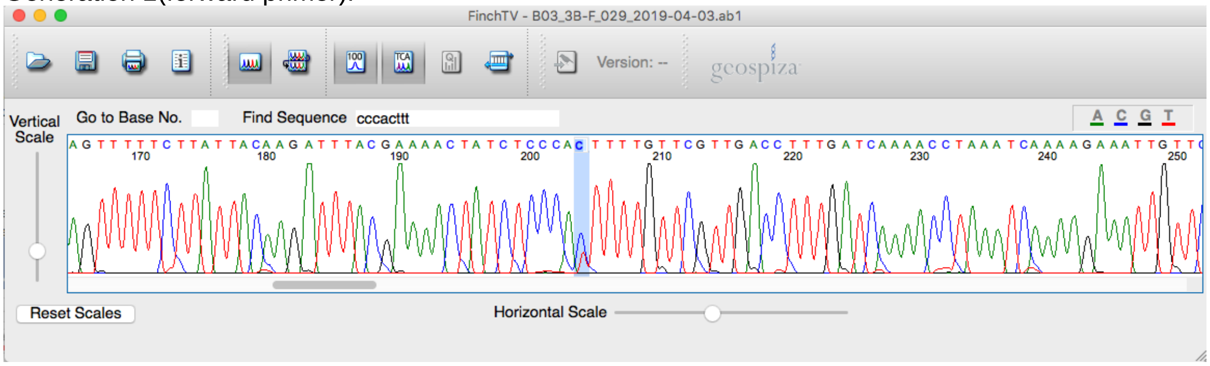


Generation 1(reverse primer):

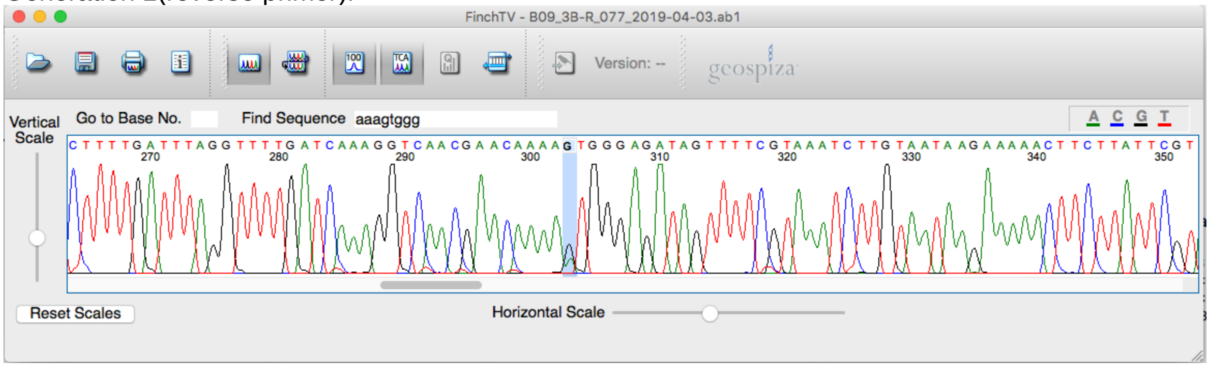

**Figure S2 (continued).** Line: 38a; Scaffold786:78964; T->T/A – High confidence – Confirmed

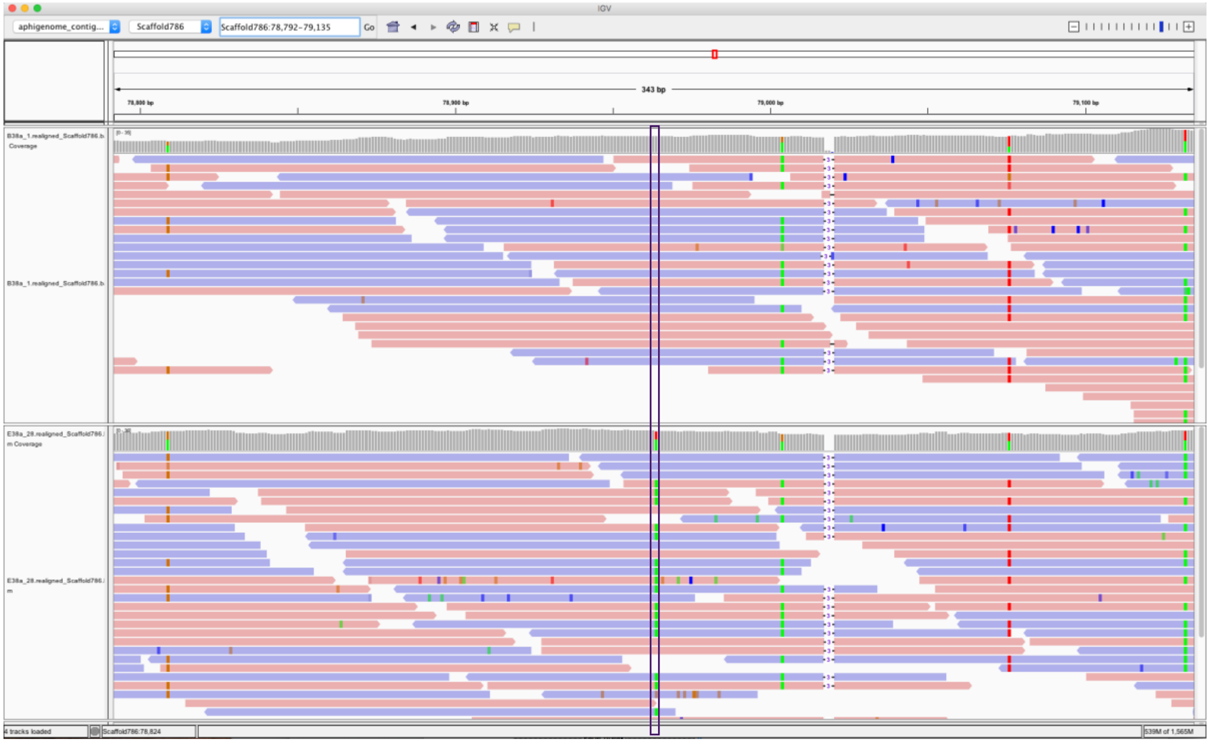


Generation 1:

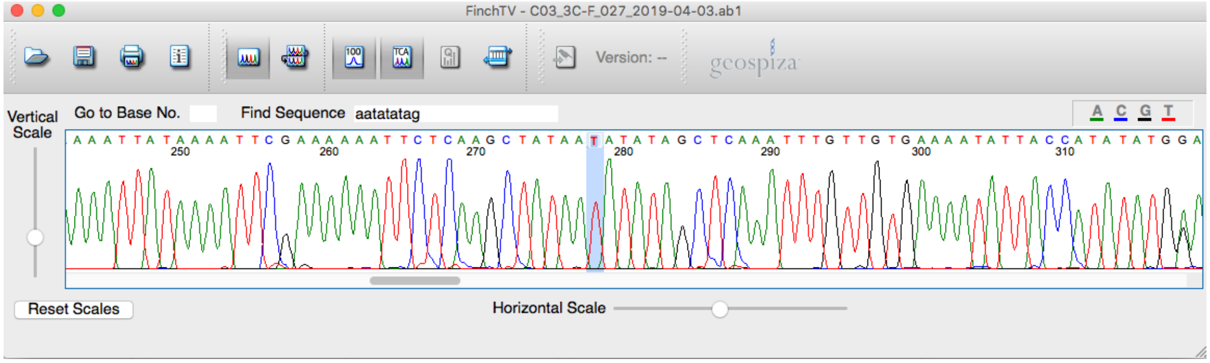


Generation 28:

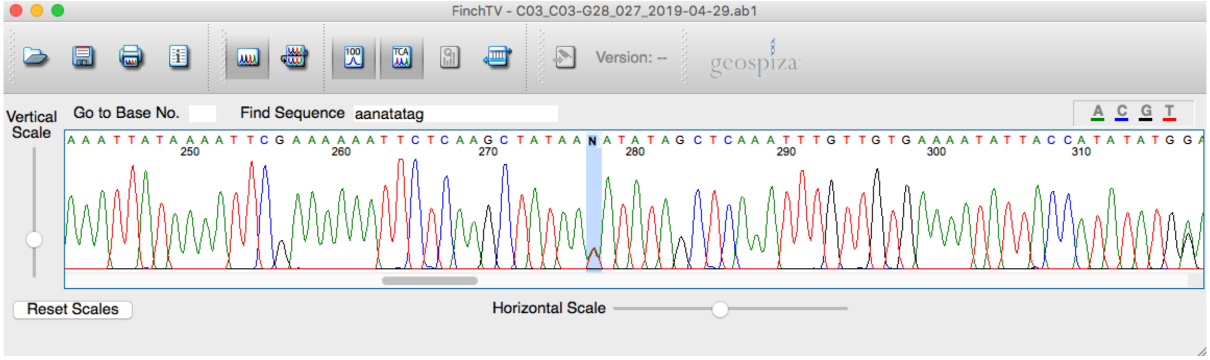

**Figure S2 (continued)**. Line 38a; Scaffold78:1958943; AR ->A/G – High confidence – Confirmed

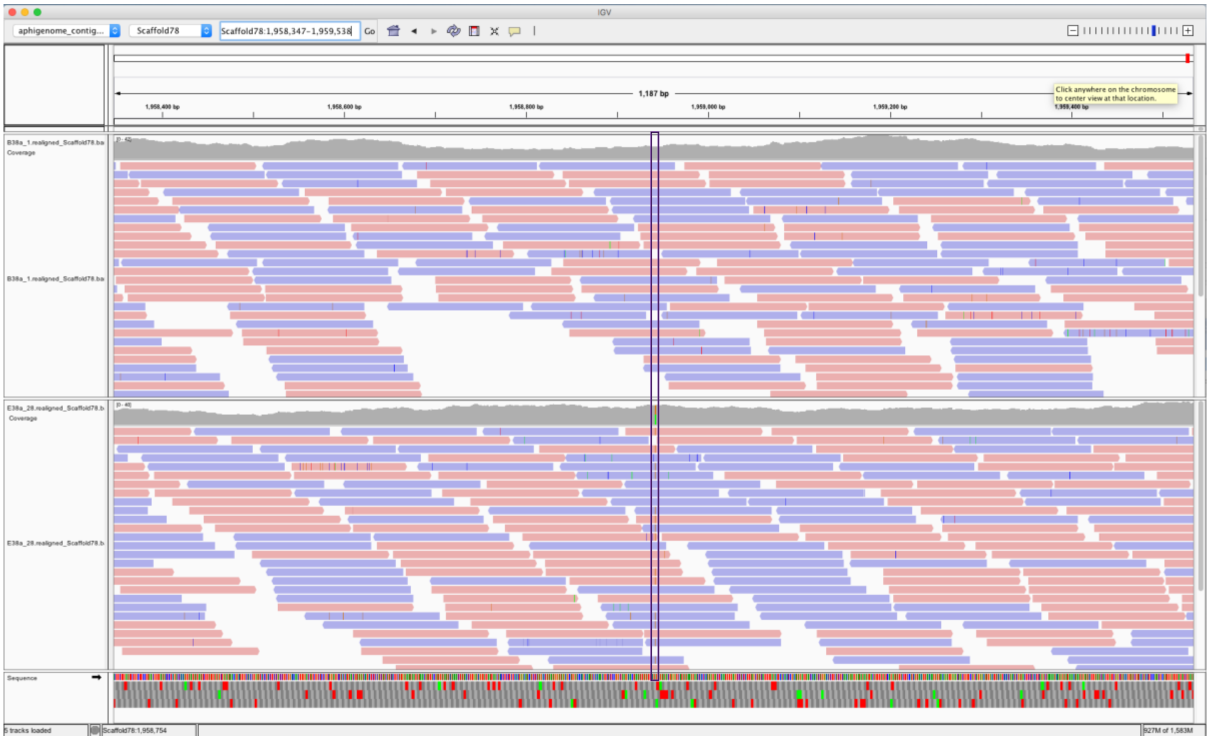

Generation 1:

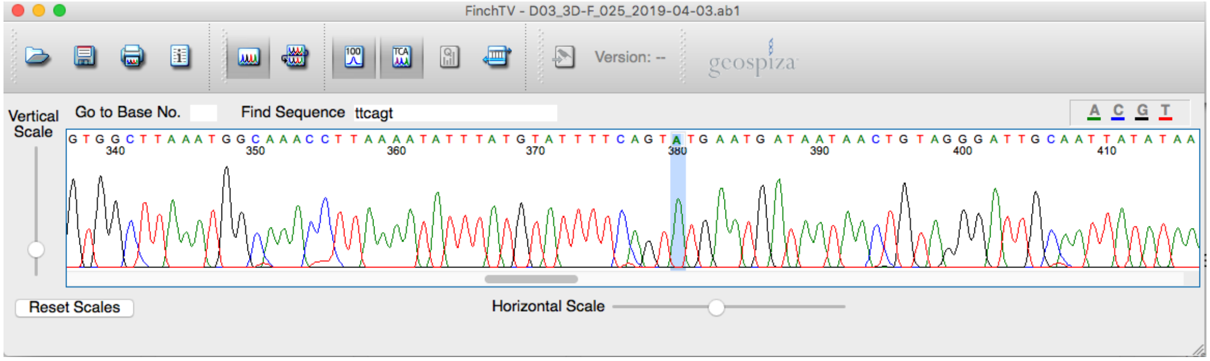


Generation 28:

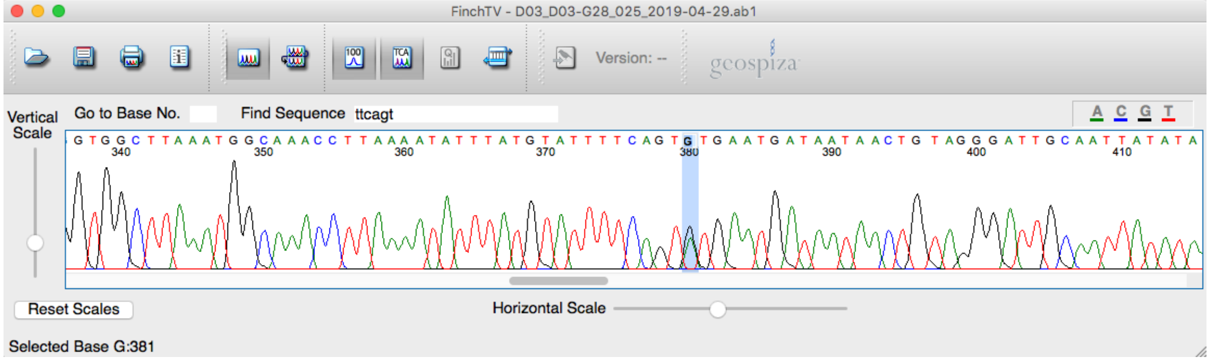

**Figure S2 (continued).** Line 54a; Scaffold149:198799; C->C/T – High confidence – Confirmed

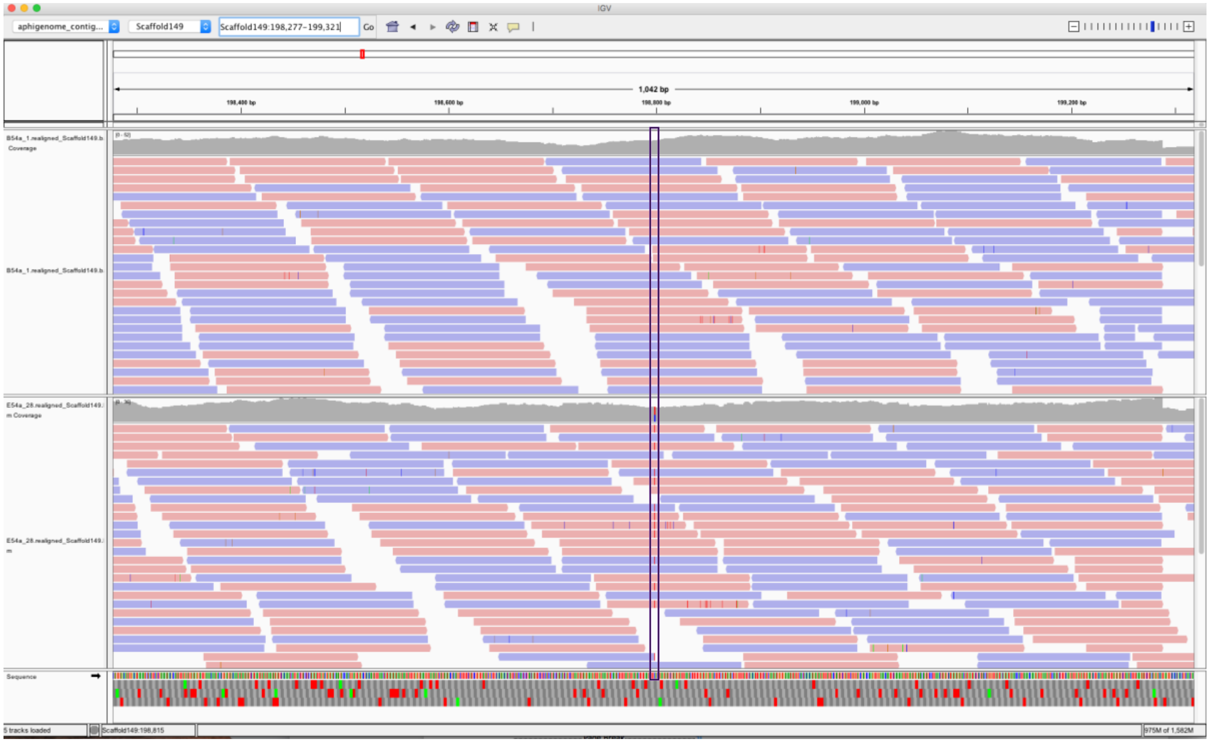


Generation 1:

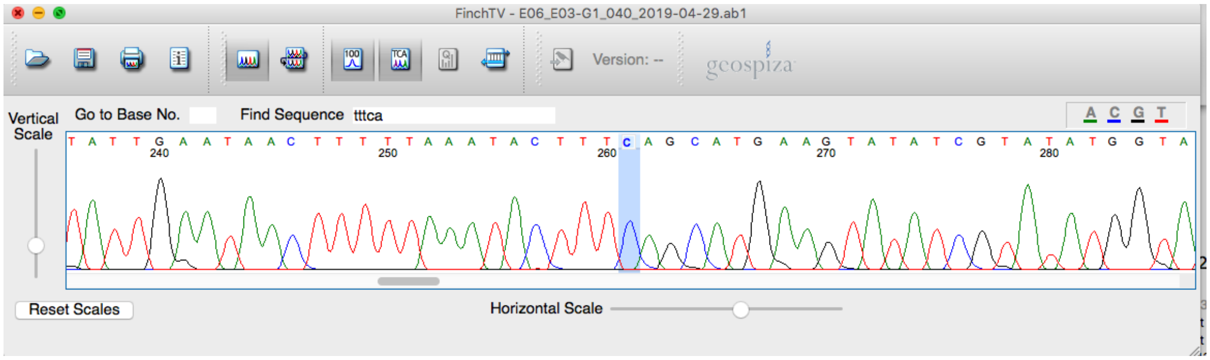


Generation 28:

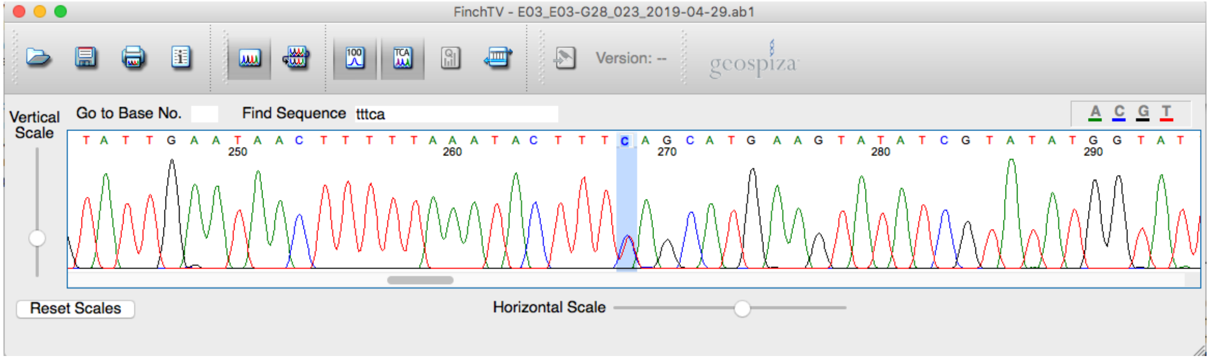

Line: 16.37a; Scaffold133: 216490; C->C/T– Low confidence (reads with T have deletions) – Not confirmed

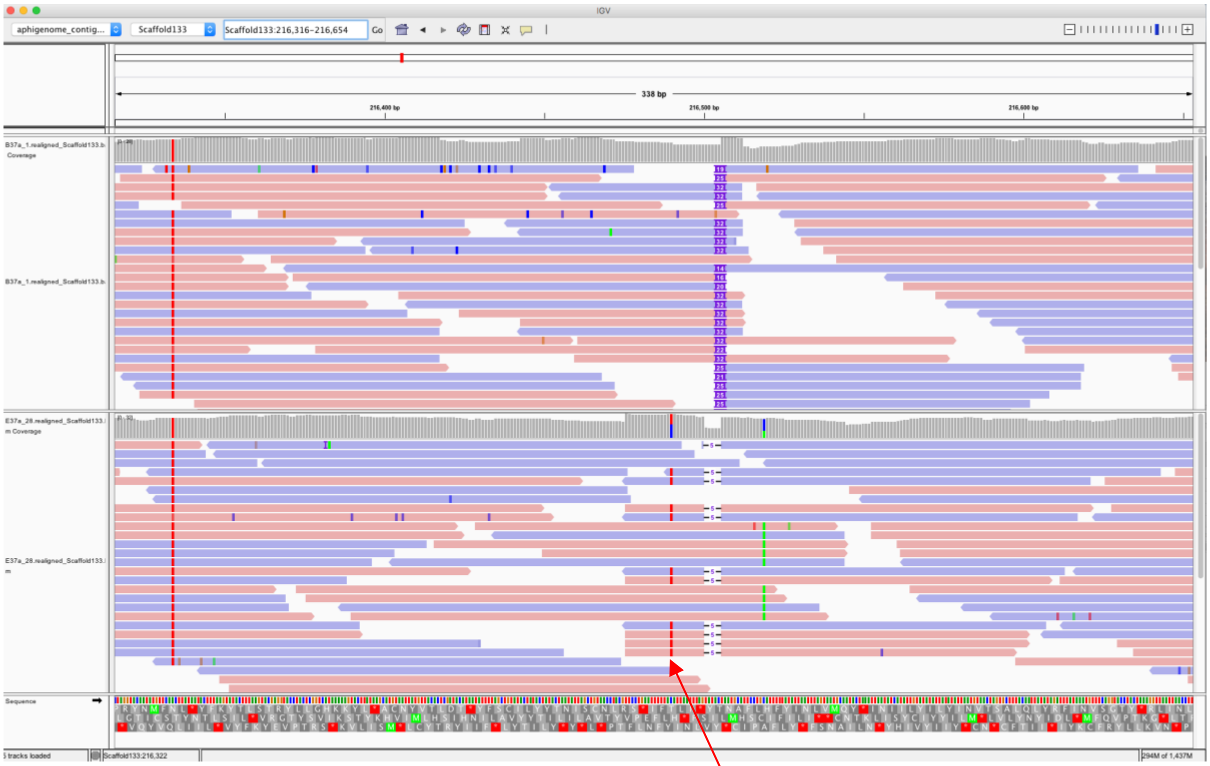


Generation 1:

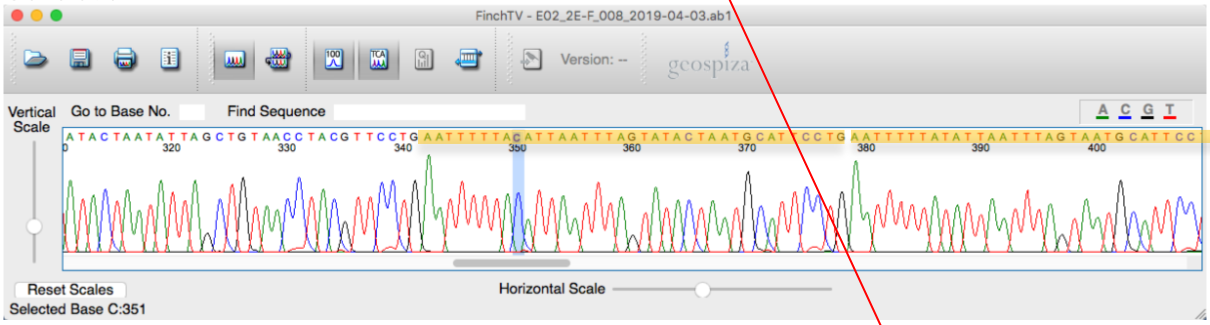


Generation 28:

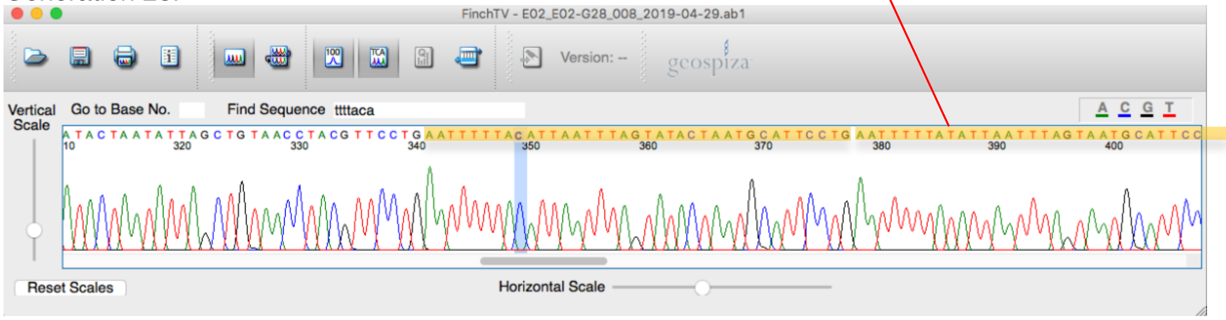


Comments: Read mapping errors in Generation 28 (IGV, bottom track). Sanger sequencing shows two repeats (highlighted by yellow rectangles), the reference genome includes only the upstream repeat. Base T of the candidate mutation in Generation 28 results from mapping of the downstream region to the upstream one. The correct base for both generations is C (highlighted by blue colour).

**Figure S2 (continued).** Line: 81a; Scaffold152:558607; C/T->C – Low confidence (heterozygote->homozygote) – Not confirmed

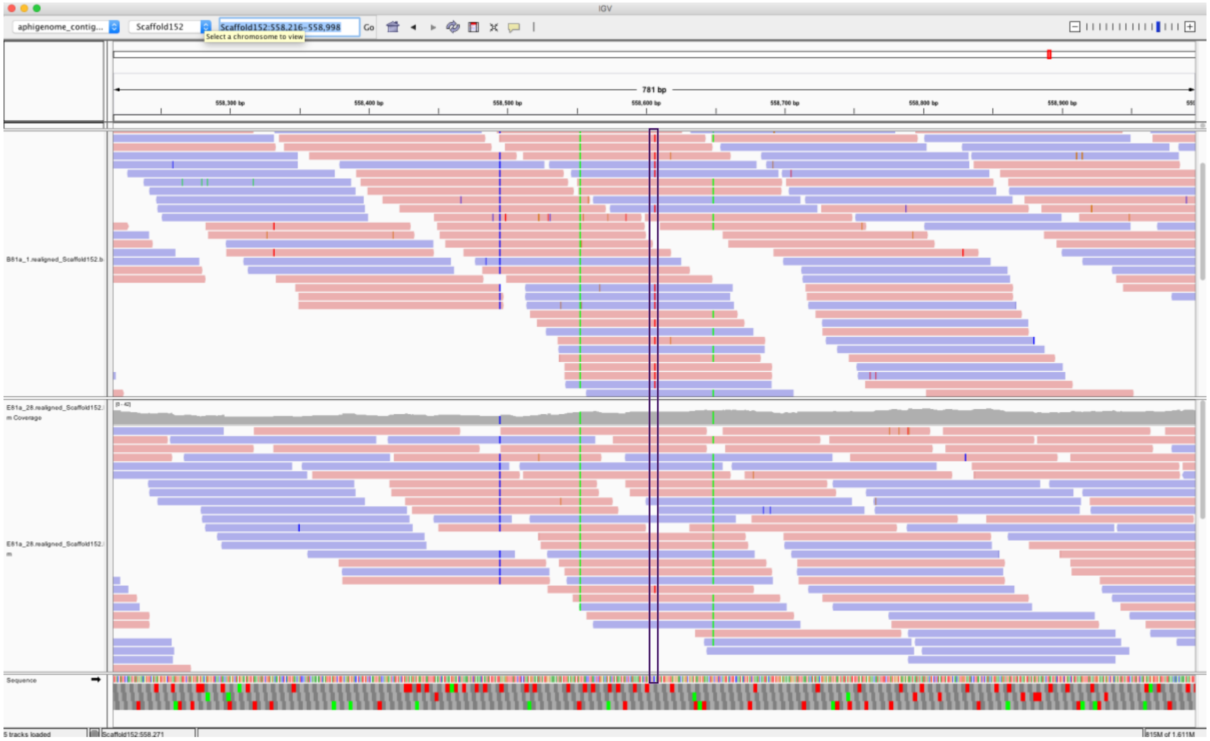

Generation 1:

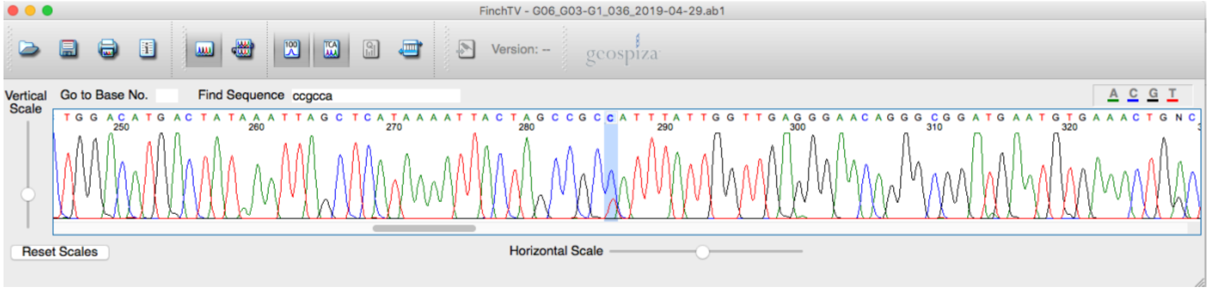

Generation 28:

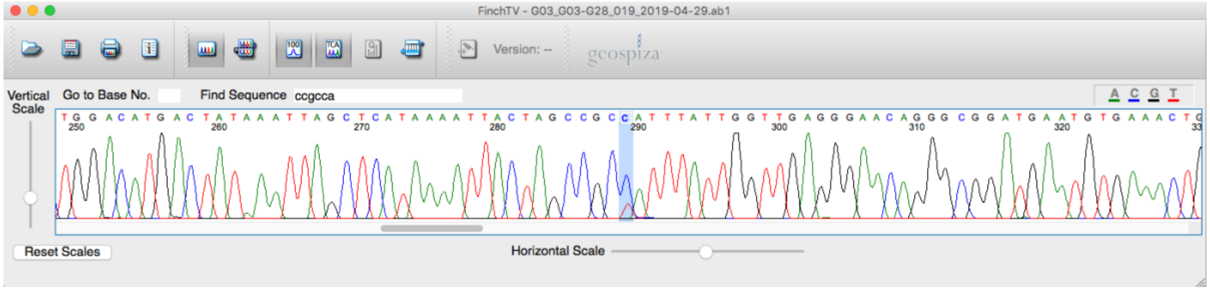

Comment: Generation 28 is in fact heterozygote (C/T).

**Figure S2 (continued).** Line: 38a; Scaffold238:90257; A->A/G Low confidence (multiple candidate mutations within 50 bp) – Not confirmed

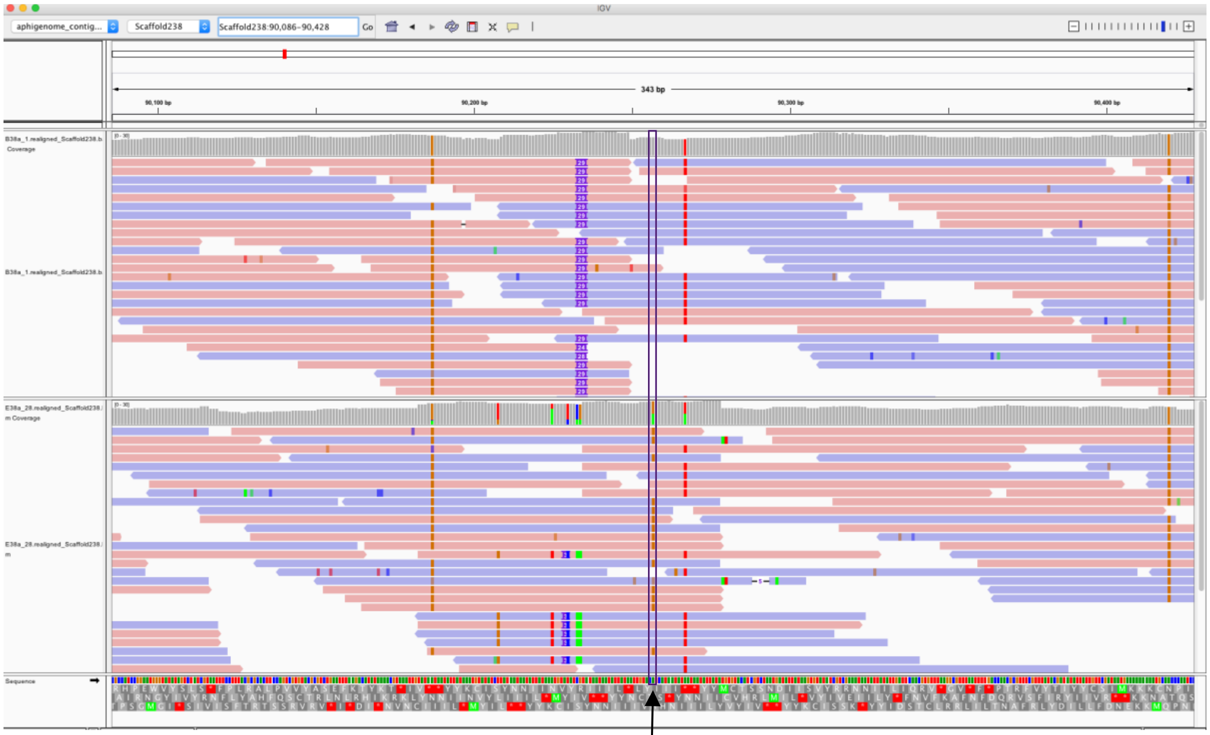


Generation 1 (sequence highlighted by blue is not present in the reference genome assembly):

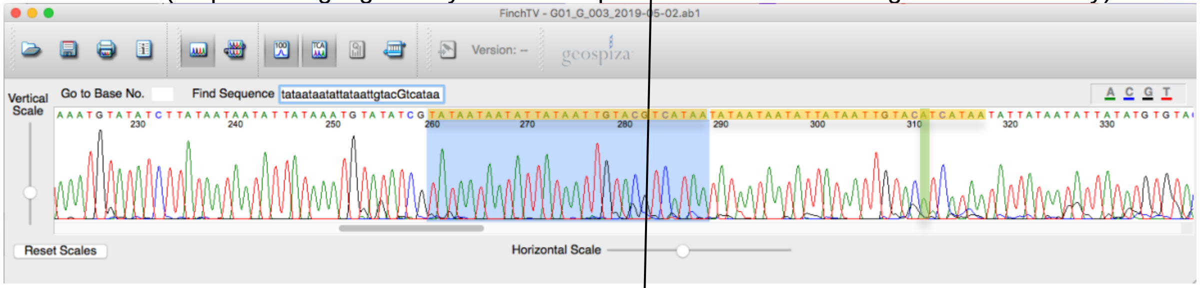


Generation 28 (sequence highlighted by blue is not present in the reference genome assembly):

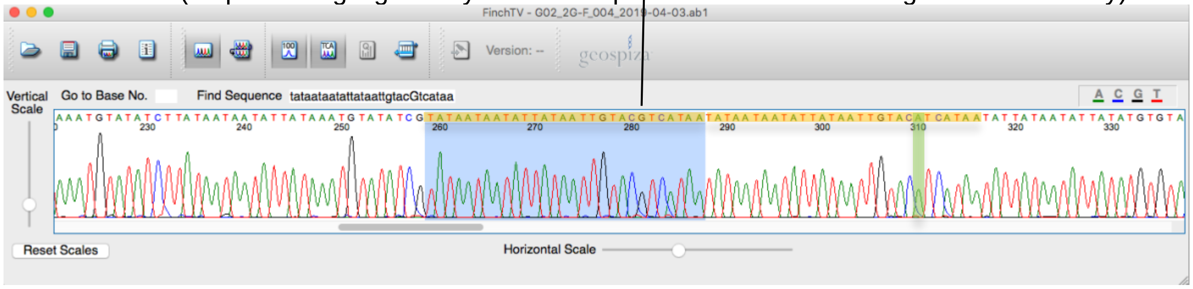


Comments: Read mapping errors in Generation 28 (IGV, bottom track). Base G of the candidate mutation (position 90257) results from mapping of the upstream region to the downstream one. The correct base for both generations is A (highlighted by green colour).

**Figure S2 (continued).** Line: PT1; Scaffold415:393952; C->C/T – Low confidence (1 read with T in Generation 1) – Not confirmed

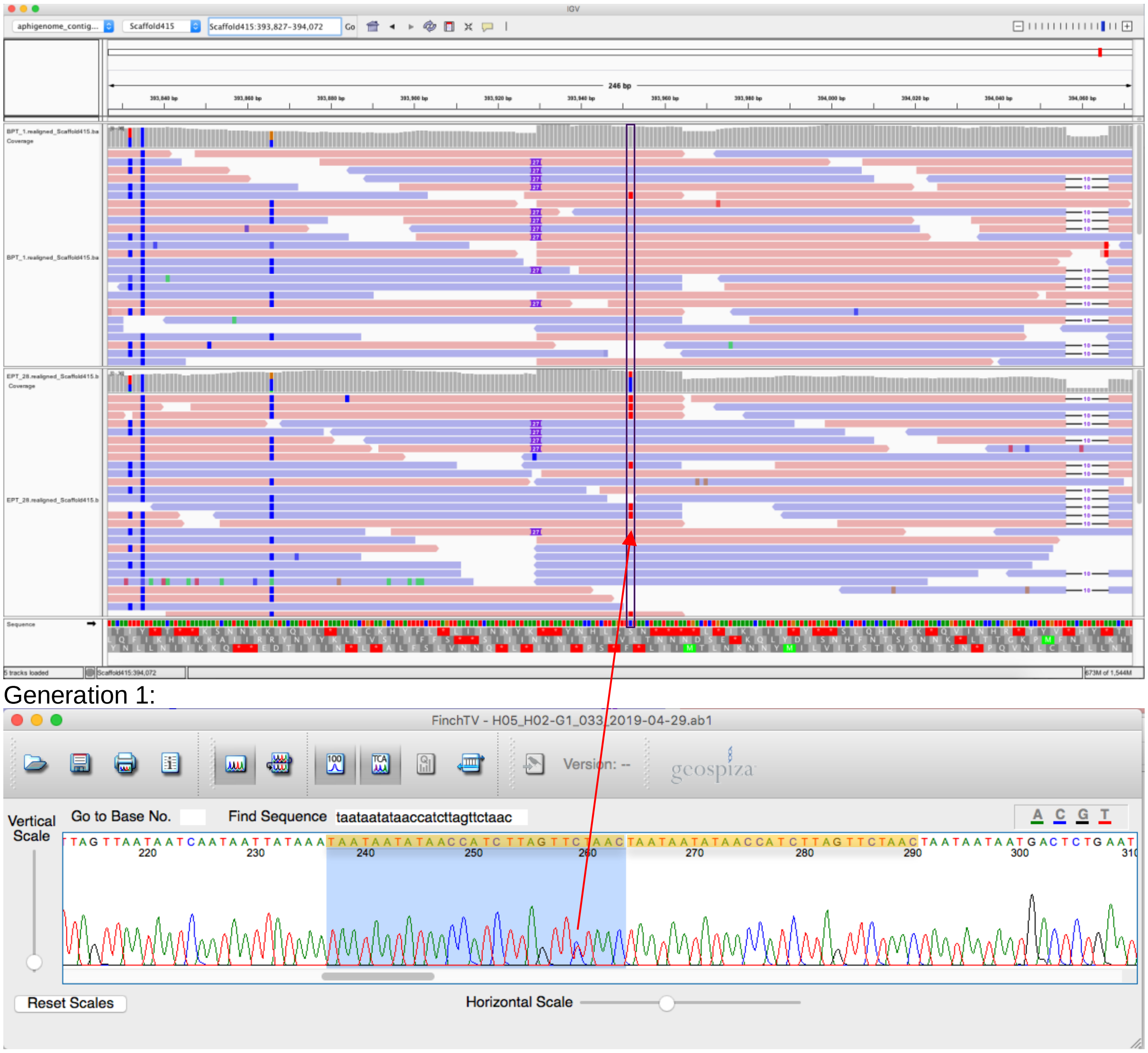


Comment: Sanger sequencing shows two repeats (highlighted by yellow rectangles), the reference genome does not have the upstream repeat (highlighted by blue rectangle). Base T of the candidate mutation results from mapping of the upstream region to the downstream one.

### Table S3. Sanger sequencing validation summary

| Line | Scaffold | Position | Forward_Primer | Reverse_Primer | PCR_result | Mutation | Generation_1_F_Primer | Generation_1_R_Primer | Generation_28_F_Primer | Confirmed |
|---|---|---|---|---|---|---|---|---|---|---|
| 05b | Scaffold59 | 515379 | GAAGAGAAGTACCGCCCGAG | TGTCCTTAAAGGCAGGGCTG | sharp band | AR -- High Confidence | A | Sanger failed | A/G | Yes |
| 09b | Scaffold153 | 382214 | TTGGCGTCACCCGTTATGTG | AAACCTTCCTGCGAACAACG | sharp band | TK -- High Confidence | T/G | Sanger failed | Sanger failed | No |
| 09b | Scaffold41 | 243055 | GGCTGTCAGGGAATCTTCTCG | AGTGTTCTTGCAAACGATCCAG | sharp band | TW -- High Confidence | T | T | T/A | Yes |
| 09b | Scaffold429 | 500010 | CGGTCATGCCACGAACTAAG | TCACAATAGCCTACTAGCGACTG | sharp band | GR -- High Confidence | G | Sanger failed | Sanger failed 2x | – |
| 16.107a | Scaffold1165 | 78180 | CAACCAGTTTGATACGCCGG | AGTGGGTTGATAATGTAAAGGTACC | weak band | YT -- Low Confidence | wrong mapping | Sanger failed | Sanger failed | No |
| 16.107a | Scaffold1165 | 78222 | CAACCAGTTTGATACGCCGG | AGTGGGTTGATAATGTAAAGGTACC | weak band | WT -- Low Confidence | wrong mapping | Sanger failed | Sanger failed | No |
| 16.107a | Scaffold688 | 78705 | GTCAAAGTGATTACCGAGTGGG | GTGGCCCTAACTTATCAATCGTC | sharp band | CY -- High Confidence | Sanger failed 2x | Sanger failed | Sanger failed | – |
| 16.118a | Scaffold330 | 305873 | TCGTACGTTCACATGTTTGGG | GTATGTACGGCAAATTGAGTAACTC | sharp band | GK -- High Confidence | Sanger failed 2x | Sanger failed | Sanger failed | – |
| 16.120a | Scaffold1559 | 38127 | TGTCTCGGCTTCAACTACGC | GCCAATGACGTCTGTATGGG | sharp band | CY -- High Confidence | C | C | C/T | Yes |
| 16.120a | Scaffold19 | 179756 | GGTACTGTGTCGTAGGACCG | ACAATACCACGGGTATACGCG | sharp band | CM -- High Confidence | Sanger failed | C | Sanger failed 2x | – |
| 16.122a | Scaffold25 | 659741 | TCGTAGCATATTCCAACAAGTCTTG | ACGTGTCATTACGAAGTGCG | sharp band | GR -- High Confidence | G | G | A/G | Yes |
| 29d | Scaffold19 | 1669887 | GAGACCCTCAAAGTCCTGCC | TCGTTGCACCTCTTTGGATC | no band | YC -- Low Confidence | | | | – |
| 29d | Scaffold20 | 513622 | AAAGTCCACACGGGTCTCTG | CCAGGATAGAAGTGTCAGTGGG | sharp band | CY -- High Confidence | C | C | C/T | Yes |
| 16.37a | Scaffold133 | 216490 | GGTATTGTGCCCACTTCACAC | ATGTAGCTCCAAGTGCACAG | sharp band | CY -- Low Confidence | C | C | C | No |
| 16.37a | Scaffold68 | 678872 | ACCAGCAAACCAGCATGATG | CAGTATGAATTTGTAGGCCAACTAC | sharp band | GR -- High Confidence | G | Sanger failed | A/G | Yes |
| 16.38a | Scaffold238 | 90257 | TGCATCTACTCGATCGCAGC | ACCCAAGTAGCCAGGCAATG | sharp band | AR -- Low Confidence | wrong mapping | wrong mapping | wrong mapping | No |
| 16.38a | Scaffold359 | 316424 | ACAACATCTTCTGCATAGGGC | GGTTTGGCTGTTGGTCAAGAG | sharp band | CM -- High Confidence | C | C | C/A | Yes |
| 16.38a | Scaffold3 | 1728624 | ATTCGTGAGTGACTCCCGTG | GAACTGATTTCTTCCGGCGC | sharp band | GR -- High Confidence | Sanger failed 2x | Sanger failed | Sanger failed | – |
| 16.38a | Scaffold42 | 194414 | TCACATCATTAGACCTCTTCCTTC | AATGTCTCGTCGAACCCTGG | sharp band | CY -- High Confidence | C/T | C/T | | No |
| 16.38a | Scaffold786 | 78964 | TGTTCATGGTAGACAATTTGTGGAG | ACCCATGAATTTGCAGCTTCAC | sharp band | TW -- High Confidence | T | Sanger failed | A/T | Yes |
| 16.38a | Scaffold78 | 1958943 | TCGCTAGCTCTTGTTCGTGG | TGTCTCACTCGACTAGGACAAC | sharp band | AR -- High Confidence | A | A | A/G | Yes |
| 54a | Scaffold149 | 198799 | TGACCCAACATGTAAGTTAGTACC | CACAAACCTTTGGGCTTCGG | double band | CY -- High Confidence | C | Sanger failed | C/T | Yes |
| 54a | Scaffold150 | 629228 | ACGATACTCGAACATTTGTACTTCC | ACAATCGATATACGTTGGCCAAC | no band | SG -- Low Confidence | | | | – |
| 16.81a | Scaffold152 | 558607 | TGGGTTCAACTAACATGAGGC | GAGCTTAAGGTATTACCAATGTTGC | sharp band | YC -- Low Confidence | C/T | Sanger failed | C/T | No |
| 16.81a | Scaffold265 | 763936 | TTGGGCATGGGTTTCCCTAC | TATTCGATAAGCCGTGCACG | sharp band | CM -- High Confidence | Sanger failed 2x | Sanger failed | Sanger failed | – |
| PT1 | Scaffold415 | 393952 | ACCTGATTTCACCCGGTAGTC | GTTGGCACCGAGCTATTCTG | sharp band | CY -- Low Confidence | wrong mapping | Sanger failed | wrong mapping | No |

**Table S4. List of high confidence mutations**

| Line | Scaffold | Position | Substitution | Mutation | Substitution type | Chromosome |
|---|---|---|---|---|---|---|
| 05b | Scaffold25 | 833375 | GR | A/G | Transition | X |
| 05b | Scaffold59 | 515379 | AR | A/G | Transition | A |
| 05b | Scaffold321 | 320441 | AR | A/G | Transition | A |
| 05b | Scaffold985 | 29540 | GR | A/G | Transition | A |
| 09b | Scaffold41 | 243055 | TW | A/T | Transversion | A |
| 09b | Scaffold124 | 40313 | CY | C/T | Transition | A |
| 09b | Scaffold153 | 382214 | TK | T/G | Transversion | X |
| 09b | Scaffold277 | 372547 | GR | A/G | Transition | A |
| 09b | Scaffold429 | 500010 | GR | A/G | Transition | A |
| 107a | Scaffold128 | 546231 | TY | C/T | Transition | A |
| 107a | Scaffold688 | 78705 | CY | C/T | Transition | X |
| 118a | Scaffold330 | 305873 | GK | T/G | Transversion | A |
| 118a | Scaffold412 | 511526 | TK | T/G | Transversion | X |
| 118a | Scaffold470 | 50266 | GR | A/G | Transition | X |
| 118a | Scaffold735 | 101714 | GR | A/G | Transition | A |
| 120a | Scaffold19 | 179756 | CM | A/C | Transversion | A |
| 120a | Scaffold63 | 876430 | CY | C/T | Transition | A |
| 120a | Scaffold134 | 189526 | TY | C/T | Transition | A |
| 120a | Scaffold246 | 236799 | TW | A/T | Transversion | A |
| 120a | Scaffold719 | 79702 | AR | A/G | Transition | A |
| 120a | Scaffold827 | 144490 | AW | A/T | Transversion | A |
| 120a | Scaffold963 | 159285 | GR | A/G | Transition | X |
| 120a | Scaffold1559 | 38127 | CY | C/T | Transition | A |
| 122a | Scaffold25 | 659741 | GR | A/G | Transition | X |
| 29d | Scaffold20 | 513622 | CY | C/T | Transition | A |
| 29d | Scaffold1083 | 207254 | GR | A/G | Transition | A |
| 37a | Scaffold10 | 1560013 | AM | A/C | Transversion | X |
| 37a | Scaffold68 | 678872 | GR | A/G | Transition | A |
| 37a | Scaffold531 | 189813 | TW | A/T | Transversion | A |
| 38a | Scaffold3 | 1728624 | GR | A/G | Transition | A |
| 38a | Scaffold4 | 1511577 | GR | A/G | Transition | A |
| 38a | Scaffold42 | 194414 | CY | C/T | Transition | A |
| 38a | Scaffold78 | 1958943 | AR | A/G | Transition | A |
| 38a | Scaffold359 | 316424 | CM | A/C | Transversion | A |

**Table S4 (continued).**

| Line | Scaffold | Position | Substitution | Mutation | Substitution type | Chromosome |
|---|---|---|---|---|---|---|
| 38a | Scaffold786 | 78964 | TW | A/T | Transversion | X |
| 54a | Scaffold149 | 198799 | CY | C/T | Transition | A |
| 54a | Scaffold502 | 146192 | AW | A/T | Transversion | A |
| 54a | Scaffold999 | 50104 | GR | A/G | Transition | A |
| 54a | Scaffold2061 | 36937 | CM | A/C | Transversion | A |
| 81a | Scaffold265 | 763936 | CM | A/C | Transversion | X |
| 81a | Scaffold310 | 181316 | GR | A/G | Transition | A |
| PT | Scaffold258 | 488981 | TW | A/T | Transversion | NA |
| PT | Scaffold416 | 377784 | CY | C/T | Transition | X |

**Table S5. Codes of pea aphid lineages and sampling locations**

| Code | Genotype | Latitude | Longitude | Collection date | Note |
|---|---|---|---|---|---|
| 05b | *Medicago lupulina* | 51.758323 | -1.296352 | 07/06/2015 | MA |
| 09b | *Lotus corniculatus* | 51.754401 | -1.278131 | 14/06/2015 | MA |
| 16.107a | *Lathyrus pratensis* | 51.761003 | -1.23489 | 15/08/2016 | both |
| 16.118a | *Lotus corniculatus* | 51.742694 | -1.194073 | 17/08/2016 | MA |
| 16.120a | *Lathyrus pratensis* | 51.743372 | -1.193655 | 17/08/2016 | both |
| 16.122a | *Lotus corniculatus* | 51.743777 | -1.194975 | 17/08/2016 | MA |
| 29d | *Lathyrus pratensis* | 51.776694 | -1.348532 | 04/07/2015 | both |
| 16.37a | *Lotus pedunculatus* | 51.68356476 | -1.201154 | 30/07/2016 | MA |
| 16.38a | *Lotus pedunculatus* | 51.68353524 | -1.201539 | 30/07/2016 | MA |
| 54a | *Medicago lupulina* | 51.747583 | -1.283362 | 04/08/2015 | MA |
| 16.81a | *Lotus pedunculatus* | 51.684389 | -1.200583 | 11/08/2016 | MA |
| PT1 | *Lotus arenaruis?* | 37.59804 | -8.816121 | 04/05/2017 | MA |
| 16.19a | *Lathyrus pratensis* | 51.730344 | -1.272286 | 02/07/2016 | PopGen |
| 16.26a | *Lathyrus pratensis* | 51.741111 | -1.250278 | 08/07/2016 | PopGen |
| 16.62a | *Lathyrus pratensis* | 51.763611 | -1.250278 | 08/08/2016 | PopGen |
| 16.80a | *Lathyrus pratensis* | 51.683611 | -1.201111 | 11/08/2016 | PopGen |
| 16.110a | *Lathyrus pratensis* | 51.770833 | -1.335278 | 16/08/2016 | PopGen |
| 16.116a | *Lathyrus pratensis* | 51.770833 | -1.335278 | 16/08/2016 | PopGen |
| 16.138 | *Lathyrus pratensis* | 51.745203 | -1.262684 | 18/08/2016 | PopGen |
| 16.161b | *Lathyrus pratensis* | 51.788798 | -1.258795 | 04/09/2016 | PopGen |
| 61b | *Lathyrus pratensis* | 51.812304 | -1.173401 | 09/08/2015 | PopGen |
| 70a | *Lathyrus pratensis* | 51.810836 | -1.175696 | 29/08/2015 | PopGen |
| 16.28a | *Vicia cracca* | 51.759867 | -1.243324 | 23/07/2016 | PopGen |
| 16.61a | *Vicia cracca* | 51.76361111 | -1.25 | 08/08/2016 | PopGen |
| 16.63a | *Vicia cracca* | 51.76583333 | -1.25056 | 08/08/2016 | PopGen |
| 16.84a | *Vicia cracca* | 51.80944444 | -1.17556 | 12/08/2016 | PopGen |
| 16.95a | *Vicia cracca* | 51.81222222 | -1.18139 | 12/08/2016 | PopGen |
| 16.105a | *Vicia cracca* | 51.761003 | -1.23489 | 15/08/2016 | PopGen |
| 16.131a | *Vicia cracca* | 51.75833333 | -1.27472 | 18/08/2016 | PopGen |
| 642a | *Vicia cracca* | 51.771147 | -1.252772 | 05/07/2013 | PopGen |
| 56b | *Vicia cracca* | 51.745872 | -1.288225 | 04/08/2016 | PopGen |
| 60c | *Vicia cracca* | 51.812357 | -1.172889 | 09/08/2016 | PopGen |
| 65a | *Vicia cracca* | 51.8104 | -1.17592 | 09/08/2016 | PopGen |
| 67a | *Vicia cracca* | 51.8123 | -1.17885 | 09/08/2016 | PopGen |

**Table S6. Software versions**

| Software | PopGen | Mutation Accumulation |
|---|---|---|
| fastqc | 0.11.3 | 0.11.8 |
| multiqc | 1.3 | 1.6 |
| trim_galore | 0.4.0 | 0.5.0 |
| cutadapt | 1.8.3 | 1.8.3 |
| bwa mem | 0.7.17-r1188 | 0.7.17-r1188 |
| GATK | 3.4-46 | 3.8-0-ge9d806836 |
| Samtools/bcftools | 1.6 (using htslib 1.6) | 1.3 (using htslib 1.3.1) |
| Picard AddOrReplaceReadGroups | 2.18.15-SNAPSHOT | 2.18.15-SNAPSHOT |
| Integrative Genomics Viewer | 2.4.7 | 2.4.7 |

**Table S7. Number of callable sites and candidate mutations for each asexual lineage at each filtering step**

| Line | Number of callable sites | Number of putative mutations (Set 1) | Bad | Triallelic | Pass (Set 2) | High_Conf (Set 3) | Low_Conf (Set 3) |
|---|---|---|---|---|---|---|---|
| 05b | 175,281,226 | 1,224 | 1,006 | 184 | 34 | 4 | 0 |
| 09b | 185,010,536 | 1,160 | 959 | 181 | 20 | 5 | 5 |
| 16.107a | 196,589,460 | 2,137 | 1,742 | 365 | 30 | 2 | 2 |
| 16.118a | 280,177,299 | 1,371 | 1,058 | 176 | 137 | 4 | 2 |
| 16.120a | 189,053,054 | 2,118 | 1,690 | 372 | 56 | 8 | 1 |
| 16.122a | 258,621,867 | 1,170 | 983 | 174 | 13 | 1 | 1 |
| 29d | 155,191,035 | 2,188 | 1,809 | 347 | 32 | 2 | 2 |
| 16.37a | 166,587,314 | 980 | 810 | 155 | 15 | 3 | 1 |
| 16.38a | 185,499,166 | 1,215 | 1,028 | 169 | 18 | 6 | 2 |
| 54a | 258,825,042 | 1,183 | 953 | 212 | 18 | 4 | 5 |
| 16.81a | 204,485,061 | 1,173 | 1,001 | 151 | 21 | 2 | 1 |
| PT1 | 213,058,132 | 1,141 | 919 | 205 | 17 | 2 | 2 |
| Total: | 2,468,379,192 | | | | 411 | 43 | 24 |

**Table S8. PCR conditions.**

| Amount | Component | Final Concentration |
|---|---|---|
| 34μl | Water | |
| 5μl | 10xPCR Buffer | 1x |
| 1μl | 10mM dATP | 200μM |
| 1μl | 10mM dCTP | 200μM |
| 1μl | 10mM dGTP | 200μM |
| 1μl | 10mM TTP | 200μM |
| 1μl | Jump start Taq (Sigma) | 0.05 units/ μL |
| 2.5μl | Forward primer | 0.5μM |
| 2.5μl | Reverse primer | 0.5μM |
| 1μl (min conc. 30 ng/μl) | DNA | |

25 cycles:
Initial denaturation: 94°C for 60s
Denaturation: 94°C for 30s
Annealing: 62°C for 45s;
Extension: 72°C for 60s
Final extension: 72°C for 60s
Hold: 4°C
